# A cortical–hippocampal communication undergoes rebalancing after new learning

**DOI:** 10.1101/2025.03.26.645547

**Authors:** Arron F Hall, Dong V Wang

## Abstract

The brain’s ability to consolidate a wide range of memories while maintaining their distinctiveness across experiences remains poorly understood. Sharp-wave ripples, neural oscillations that occur predominantly within CA1 of the hippocampus during immobility and sleep, have been shown to play a critical role in the consolidation process. More recently, evidence has uncovered functional heterogeneity of pyramidal neurons within distinct sublayers of CA1 that display unique properties during ripples, potentially contributing to memory specificity. Despite this, it remains unclear exactly how ripples shift the activity of CA1 neuronal populations to accommodate the consolidation of specific memories and how sublayer differences manifest. Here, we studied interactions between the anterior cingulate cortex (ACC) and CA1 neurons during ripples and discovered a reorganization of their communication following learning. Specifically, using a generalized linear model decoder, we demonstrated the pre-existence of ACC-to-CA1 communication, which is weakened during post-training sleep following learning, suggesting that ACC activity reallocates the contribution of CA1 neurons during memory formation. Interestingly, the reorganization appeared unique for a subset of CA1 superficial (CA1sup) neurons that were task inactive, whereas communication between the ACC and CA1 deep neurons remained largely stable across pre- and post-training sleep. Consistent with this sublayer-selective reorganization, we found that optogenetic stimulations of the ACC preferentially suppressed CA1sup neurons while activating a unique subset of CA1 interneurons. Overall, these findings highlight an important role of the ACC in rebalancing CA1 neuronal populations’ contribution in learning and memory consolidation.

## Introduction

Memories are not static, rather they are gradually consolidated into long-term traces across days, weeks, or longer (Klinzing et al., 2019; Squire et al., 2015). At any given moment, we are processing and consolidating an array of differing experiences. The ability to preserve distinctiveness across these various experiences is essential for memory consolidation. This requires a delicate balance between the mechanisms that facilitate the reactivation and transformation of new memories alongside those which regulate consolidation to prevent the intermingling of distinct experiences. While much attention has been focused on uncovering the principles that guide the reactivation and consolidation of memories, less is known about the mechanisms that regulate these processes and govern memory specificity.

During sleep, sharp-wave ripples, neural oscillations predominantly within the CA1 of the hippocampus, facilitate hippocampal reactivations and accompany cortical reactivations of memory-related neurons, a process referred to as replay (Buzsáki, 2015; Buzsáki et al., 1992; Davidson et al., 2009; Diba & Buzsáki, 2007; Foster & Wilson, 2006; Lee & Wilson, 2002). Replay has been shown to be essential for memory consolidation as disruption or enhancement of ripples impairs or improves memory, respectively (Ego-Stengel & Wilson, 2010; Fernández-Ruiz et al., 2019; Girardeau et al., 2009; Gridchyn et al., 2020; Jadhav et al., 2012; Wang et al., 2015). Recently, evidence has uncovered functional and anatomical heterogeneity in CA1 neurons based on their position within the pyramidal layer: the superficial (CA1sup) and deep (CA1deep) sublayers (Danielson et al., 2016; Geiller et al., 2017; Hall & Wang, 2023; Harvey et al., 2023; Mizuseki et al., 2011; Sharif et al., 2021). CA1sup neurons have more stable firing rates and higher spatial acuity (Danielson et al., 2016; Harvey et al., 2023; Mizuseki et al., 2011), whereas CA1deep neurons respond more to enriched environments, sensory landmarks, and reward (Danielson et al., 2016; Geiller et al., 2017; Harvey et al., 2023; Sharif et al., 2021). During replay events CA1sup neurons show increased activity after spatial learning, whereas CA1deep neurons show decreased activity (Berndt et al., 2023; Harvey et al., 2023). Together, ripples and the recruitment of distinct subpopulations within CA1 drive new memory formation. Despite this understanding, the mechanisms which regulate their activity to balance consolidation across experiences remain unclear.

Emerging evidence has demonstrated the importance of suppression within the hippocampal network to balance neuronal excitability surrounding learning (Gava et al., 2024; Karaba et al., 2024; Norimoto et al., 2018). During pre-learning sleep, ripples depotentiate synapses which, if perturbed, impairs learning (Norimoto et al., 2018). Similarly, after learning, ripples facilitate the depotentiation of memory-unrelated neurons (Norimoto et al., 2018). Additionally, during post-learning sleep, hippocampal neurons initially display heightened firing rates before ultimately returning to baseline (Giri et al., 2019; Wilson & McNaughton, 1994). Prior studies have demonstrated that reestablishing baseline is necessary for learning, as excessive neuronal activity and synchrony impairs learning (Gava et al., 2024; Karaba et al., 2024). Suppression of the hippocampus, then, appears to play a key role in down regulating synapses, preventing memory saturation, and promoting memory flexibility (Balduzzi & Tononi, 2013). Interestingly, CA1sup neurons appear to be the most sensitive to this process (Gava et al., 2024). Following repetitive learning, CA1sup neurons become highly synchronized and coactive, which impairs future memory formation (Gava et al., 2024). Correspondingly, optogenetic inhibition of CA1sup, but not CA1deep neurons, after learning decouples and decreases neuronal activity which reduces rigidity and restores new memory formation (Gava et al., 2024). Still, exactly how facilitation and suppression are balanced throughout consolidation, and whether cortical inputs play a role in modifying hippocampal activity in this process, remains poorly understood.

Here, we employ *in vivo* electrophysiology recording across contextual fear conditioning (CFC) and sleep to understand how communication between the anterior cingulate cortex (ACC) and hippocampal CA1 are modified following learning. Previous studies have shown that the ACC displays increased activity immediately preceding CA1 ripples and is instrumental in CFC and memory consolidation (Einarsson & Nader, 2012; Frankland et al., 2004; Wang & Ikemoto, 2016). We uncovered a novel, sublayer specific, line of communication between ACC and CA1 surrounding ripples with ACC preferentially communicating with CA1sup neurons. This communication rebalances following learning, suggesting a potential role in memory consolidation.

## Results

### Pre-ripple ACC activity predicts CA1 activity during ripples

To understand how the ACC and hippocampus communicate surrounding learning, we employed simultaneous dual-site *in vivo* electrophysiology recordings of both regions throughout a contextual fear conditioning paradigm (Figure 1A, B). In this approach, we were able to examine ACC–CA1 communication prior to, during, and after learning, enabling us to examine how this communication evolves across fear learning. Specifically, we emphasized investigation into communication changes between pre- and post-training sleep to understand whether functional connectivity undergoes learning-related reorganization. Local field potentials and neuronal spikes were recorded simultaneously, and slow-wave sleep was identified by CA1 delta-to-theta power ratio and the animal’s immobility (Figure 1C) (Wang et al., 2015).

**Figure 1.**
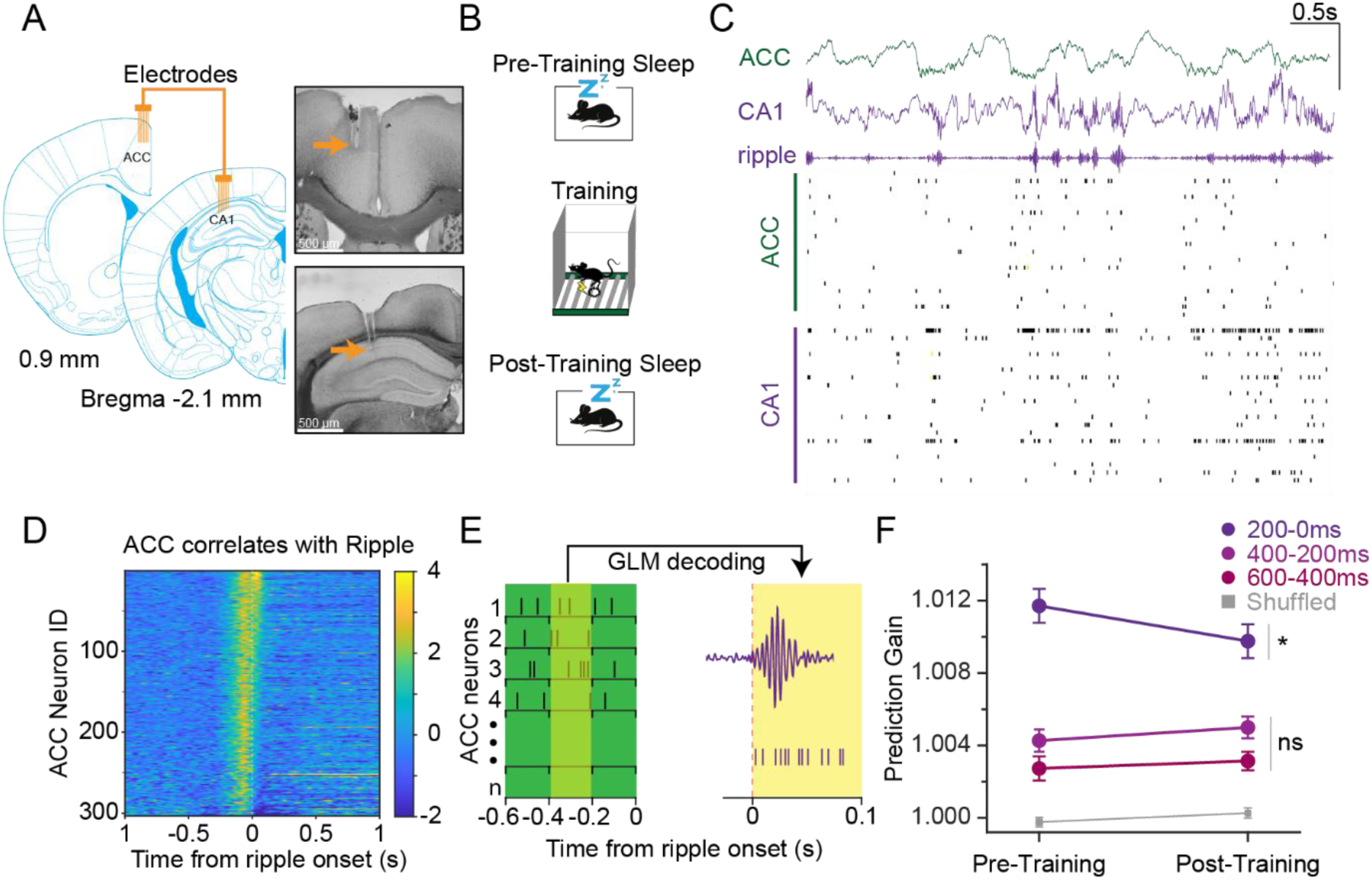
Pre-ripple ACC activity predicts CA1 activity during ripples. **A**, Left, schematic of a dual 8-tetrode array implanted in the ACC and CA1. Right, two representative brain sections highlighting the recording sites (orange arrows) in the ACC and CA1, top and bottom, respectively. **B**, Schematic of the contextual fear memory procedure. **C**, Top, Representative ACC and CA1 LFP from a pre-training sleep session, Y axis scale bar 1 mV. Bottom, individual spikes across ACC and CA1 neurons. **D**, Heatmap of ACC neuron (N = 10 animals; N = 303 neurons) activity during pre-training sleep surrounding ripple onsets (bin = 5 ms). **E**, Schematic of GLM decoder. 200 ms bins of ACC spiking data preceding ripple predict CA1 activity 0–100 ms after ripple onset. **F**, Prediction gain difference in decoding CA1 activity between real and shuffled ACC activity across three pre-ripple predictor windows (–200 to 0 ms, –400 to –200 ms, and –600 to –400 ms; N = 10 animals; N = 291 CA1 neurons). Repeated measures mixed-model analysis revealed a significant effect of predictor type, with real prediction gain exceeding shuffled prediction gain overall (*p <* .001), and a significant effect of predictor window (*p <* .001). There was also a significant effect of session (pre vs post) on prediction gain that varied across predictor windows (Window × session interaction, *p =* .010). Post-hoc comparisons showed that prediction gain was highest in the –200 to 0 ms window relative to earlier windows during pre- and post-training sleep (all Holm-corrected, *p* < .001). Additionally, the –200 to 0ms window exhibited a significant pre-to-post decrease in prediction gain (Wilcoxon signed-rank, *p* = .022), whereas the other windows did not. Error bars indicate ± SEM.

We first examined ACC neuronal spiking activity surrounding CA1 ripples during pre-training sleep. Across 10 mice, we recorded 303 ACC neurons, with individual counts per animal ranging between 19–48 neurons. Most ACC neurons (235/303) displayed significant changes in activity surrounding ripples compared to baseline (Figure 1D; see Methods). Of the neurons with significant changes, the majority displayed a pre-ripple activation (81.7%; 192/235) while a minority (18.3%; 43/235) showed a more prominent post-ripple activation (Figure 1D). To determine whether this coordinated activity is indicative of information flow, we implemented general linear model (GLM) machine learning decoding to test whether population ACC activity preceding ripples can predict CA1 activity during ripples (Figure 1E) (Jun Liu et al., 2024; Rothschild et al., 2017). ACC spiking was binned into 200 ms segments across multiple time windows to predict individual CA1 neuronal firing rates between 0–100 ms after ripple onset (Figure 1E; see Methods). The decoder was trained on 50% of the recorded data upon which the remaining 50% data was used for testing. All GLM decoding was performed on spike data recorded during slow-wave sleep epochs before or after training. We found that across multiple time-windows, prediction gain (PG) score using real data was significantly higher than shuffled (Figure 1F). PG score was greatest when using ACC spiking **–**200–0 ms preceding ripple onset, aligning with the observed correlated firing seen in Figure 1D. Moreover, only this window showed learning-related changes with a significant decrease in PG score in post-training sleep. In accordance, all subsequent GLM analyses in this paper will use the **–**200–0 ms time window (Figure 1F).

Our GLM analysis generated a PG score for each CA1 neuron, enabling us to examine how PG scores correlate and shift with other properties of CA1 neurons. We first asked whether the learning-related reduction in PG score reflected a global dampening of ACC→CA1 communication or a selective redistribution across CA1 neurons. To test this, we examined the stability of PG scores between pre- and post-training sleep. Preservation of PG score ranking among CA1 neurons would suggest that learning weakens existing ACC→CA1 communication while maintaining the relative contribution of individual ACC neurons, consistent with a global dampening effect. In contrast, poor preservation of PG score ranking would be more indicative of a reorganization of ACC→CA1 communication. We found that PG scores were positively correlated across sessions, that is, CA1 neurons with high PG scores before learning tended to retain relatively high PG score after learning (Figure 2A). To further assess the stability of PG score ranking, we performed permutation-based preservation analyses (Figure 2B). Neurons in the top PG score quartile were significantly more likely than expected by chance to remain in the top quartile following learning, whereas transitions between top and bottom quartiles were rare. Together, these findings suggest that functional connectivity between the ACC and CA1 is broadly dampened after learning rather than undergoing robust reorganization.

**Figure 2.**
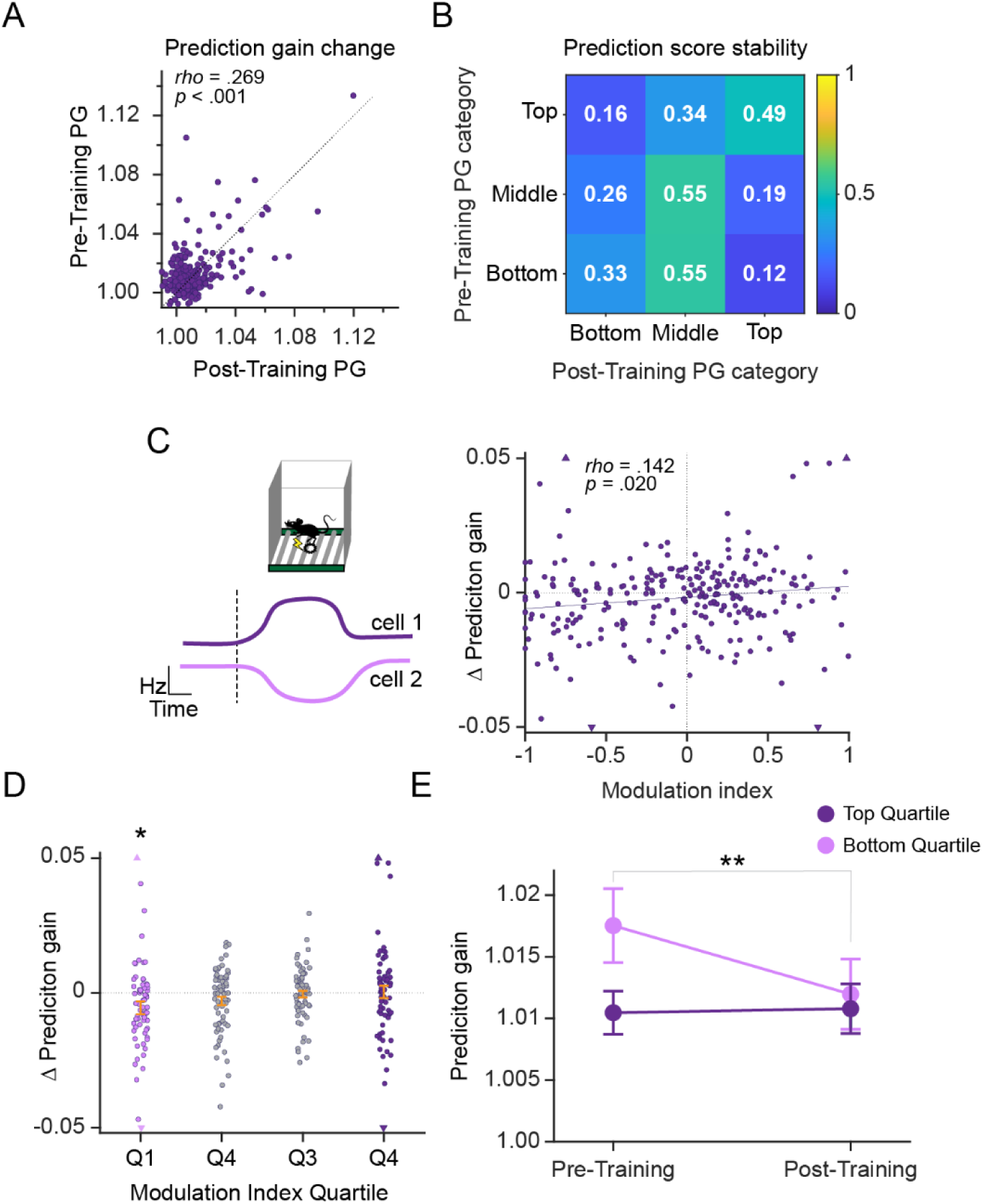
Learning dampens ACC→CA1 communication which differs based on task engagement. **A**, Correlation of PG scores between pre- and post-training sleep (N = 10 animals; N = 291 CA1 neurons). Spearman’s Rho revealed a significant correlation between pre- and post-training sleep, *rho* = .269 *p* < .001. **B**, PG score rank preservation permutation matrices. Permutation testing revealed that nearly half of CA1 neurons in the top PG score quartile remained in the top quartile after training (49.3%, permutation *p* < .001), significantly exceeding chance expectations. Whereas retention of bottom-quartile neurons was weaker, showing only a trend above chance (32.9%, permutation *p* = .052). Extreme transitions between top and bottom categories were uncommon and not enriched above chance (*p* = .999). **C**, Left, Schematic of two representative neurons’ firing patterns during training, Neuron 1 increases and Neuron 2 decreases activity during training. Right, correlation scatter plot of CA1 neuron ΔPG and modulation index. Spearman’s Rho revealed a modest positive correlation between variables, *rho* = .142, *p* = .020. Purple triangles indicate extreme values that were included in all statistical analyses but are cropped for visualization purposes. **D**, ΔPG as a function of modulation index quartiles (N = 66 per quartile). Overall quartile effect did not reach significance (Kruskal-Wallis, *p* = .081). **E**, PG scores as a function of task modulation. Neurons were divided into the top and bottom modulation index (MI) quartiles. A linear mixed-effects model revealed significant effects of MI quartile (*p* = .005) and session (*p* = .017), with a trend-level MI quartile × session interaction (*p* = .074). Post-hoc comparisons showed a significant pre-to-post reduction in PG score for neurons in the bottom MI quartile (Holm-corrected, *p* = .010), whereas neurons in the highest MI quartile showed no significant change (Holm-corrected, *p* = .831). Error bars indicate ± SEM.

To examine ACC–CA1 communication independently of the GLM analysis, we computed spike-triggered averages (STA) of CA1 neuron activity using a pooled ACC spike train generated by merging spikes from all recorded ACC neurons. We performed a full lag-resolved analysis in which CA1 spike trains were aligned to ACC spike times, baseline-normalized, and quantified in 10-ms bins across a peri-ACC-spike window (Figure 1—figure supplement 1A, B). This analysis revealed a significant reduction in CA1 activity following ACC population spiking in post-compared to pre-training sleep, suggesting a reduction in ACC–CA1 coupling after learning, consistent with the GLM decoding results.

### Task-inactive CA1 neurons reshape their activity with ACC after learning

Since overall functional connectivity between ACC and CA1 changed after learning, we next asked whether PG scores across CA1 neurons differed based on their activity during training. To test this, we calculated a modulation index by comparing each CA1 neurons’ firing rate during training with its pre-training baseline (Figure 2C) (Karaba et al., 2024). We found that predication gain change (ΔPG), showed a modest positive correlation with modulation index, indicating that task engagement modifies ACC→CA1 communication (Figure 2C). To investigate this relationship further, CA1 neurons were separated into modulation-index quartiles and their ΔPG were compared across quartiles. Although the overall effect of quartile did not reach significance in the linear mixed-effects model, neurons in the bottom quartile (task-inactive) exhibited a significant reduction in ΔPG, whereas no significant changes were observed in the remaining quartiles (Figure 2D). To better illustrate this effect, we next compared pre- and post-training PG scores in the top and bottom quartiles. Consistent with the ΔPG analysis, only neurons in the bottom quartile exhibited a significant reduction in PG scores following learning, suggesting that the that learning-related decreases in ACC→CA1 communication is primarily driven by a loss in functional connectivity with bottom quartile CA1 neurons (task-inactive) (Figure 2E). Lastly, we examined whether PG scores correlated with the freezing response in mice during recall. Across all mice, freezing was significantly higher during recall than pre-shock baseline during training (Figure 2—figure supplement 1A). Overall, we found no correlation between PG and freezing (Figure 2—figure supplement 1B–D). However, there was a trend for a positive correlation (p=.07) between ΔPG and freezing percentage. An important consideration is that the uniformly high levels of freezing in mice limited behavioral variability, potentially reducing our ability to detect relationships between ACC–CA1 communication decoding and behavioral differences.

### ACC displays a distinct relationship with CA1 sublayers

It has been well established that functional heterogeneity exists within the pyramidal layer of CA1 (Hall & Wang, 2023; Mizuseki et al., 2011). To investigate whether ACC communication is different across CA1 sublayers, we separated CA1 neurons into CA1sup and CA1deep neurons based on their sharp-wave deflection characteristics (Figure 3A; for details see Methods) (Danielson et al., 2016; Mizuseki et al., 2011). We first asked whether CA1 sublayers displayed differences in GLM decoding after learning. Although PG score was significantly higher for each sublayer compared to shuffled, neither sublayer showed a learning-related difference (trend *p* = .058) nor was there a significant difference between sublayers (Figure 3B). While GLM decoding revealed similar PG scores across CA1 sublayers, it remains possible that the stability of predictive relationship differed between them. Therefore, we examined how well PG scores correlated between pre- and post-training sleep and the degree of retention for the highly predicted neurons. Our results show that CA1sup neurons showed no significant correlation in PG scores across sessions, and permutation testing revealed that the highly predicted neurons were not retained above chance levels (Figure 3C, D). In contrast, CA1deep neurons exhibited significant PG score preservation across sessions, with both positive pre-to-post correlations and above-chance retention for the highly predicted neurons (Figure 3E, F). Together, these results suggests that ACC→CA1sup communication is more dynamic and evolving across learning, whereas ACC→CA1deep communication remains comparatively stable.

**Figure 3.**
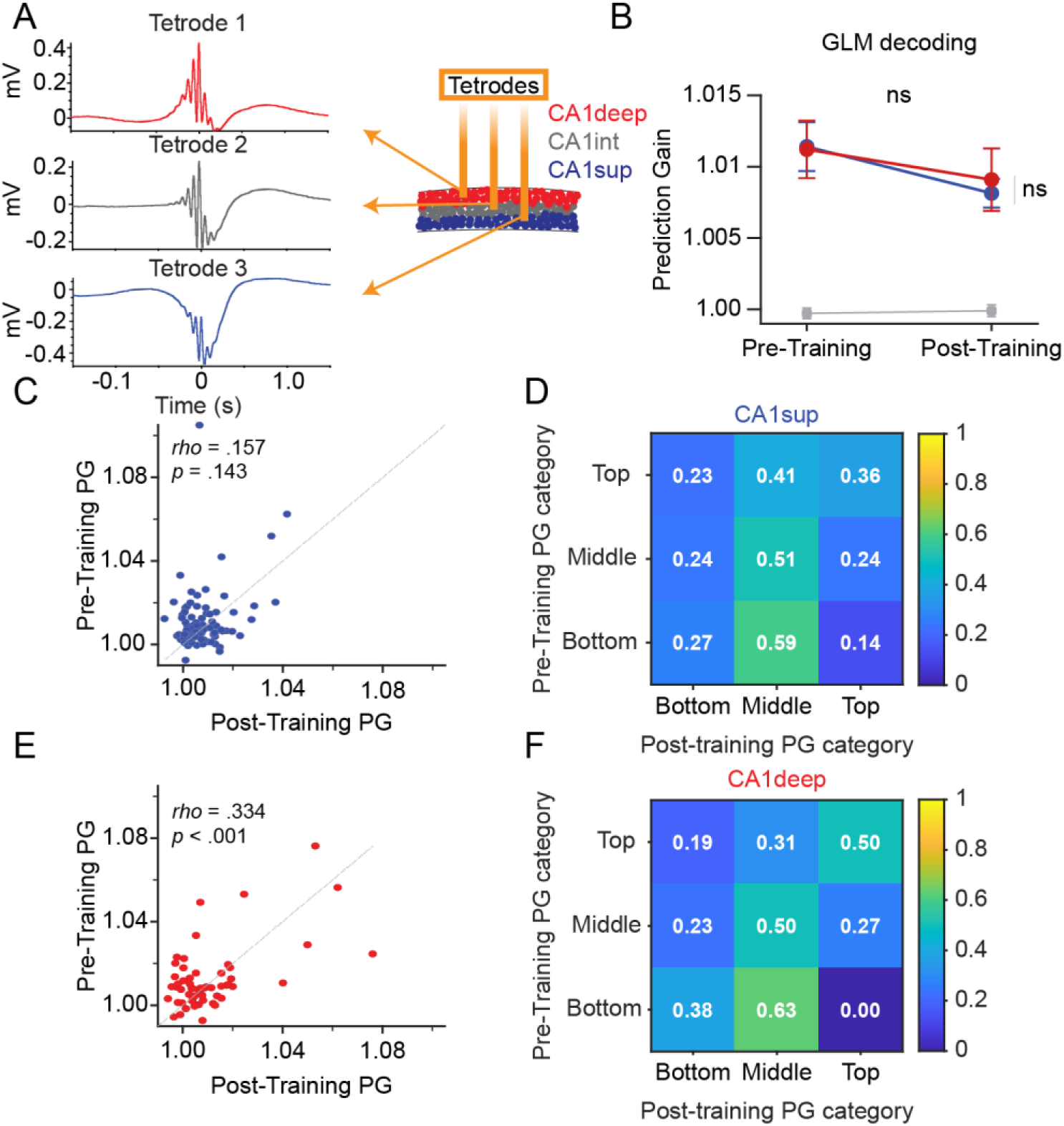
Prediction Gain stability differences between sublayers. **A**, Schematic of sublayer identification based on sharp-wave deflection difference across the pyramidal layer. **B,** PG scores before and after learning (N = 10 animals; CA1sup: N = 89; CA1deep: N = 61). Mixed-effects model revealed no significant difference in PG scores between sublayers (*p* = .755) but showed a trend for training session (*p* = .058). **C**, Correlation scatter plot of PG score and modulation index for CA1sup neurons (N = 89). Spearman’s Rho revealed no correlation between pre-training and post-training PG scores (*rho* = .157*; p* = .143). **D**, PG score rank preservation permutation matrices for CA1sup neurons (N = 89). Permutation testing revealed that neither top-quartile (36.4%, *p* = .121) nor bottom-quartile (27.3%, *p* = .484) retention exceeded chance levels. **E,** Correlation scatter plot of PG score and modulation index for CA1deep neurons (N = 61). Spearman’s Rho revealed a significant positive correlation between pre- and post-training PG scores for CA1deep neurons (*rho = .334; p* < .001). **F**, PG score rank preservation permutation matrices for CA1deep neurons (N = 61). CA1deep neurons’ top-quartile retention was 50.0% and significantly above chance, permutation *p* = .014. In contrast, bottom-quartile retention did not exceed chance levels, *p* = .174. Extreme top-to-bottom or bottom-to-top switching occurred in 8 CA1sup neurons (18.2%, *p =* .902) and 3 CA1 deep neurons (9.4% *p* = .998). Error bars indicate ± SEM.

Given the differences in PG score stability across CA1 sublayers, we next asked whether task engagement further influenced ACC→CA1 sublayer connectivity. We first examined whether ΔPG correlated with modulation index. In CA1sup neurons, we found a significant correlation between ΔPG and modulation index, suggesting that greater change in task engagement is associated with greater PG score change (Figure 4A). In support, ΔPG significantly differed across modulation index quartiles for CA1sup neurons (Figure 4B). Interestingly, only bottom quartile neurons exhibited learning-dependent change in PG with a significant reduction after learning, suggesting that the reduction in ACC→CA1 communication is mediated primarily by the loss of connectivity with task-inactive CA1sup neurons (Figure 4B, C). Together, these results suggest that CA1sup neurons that are more strongly suppressed during new learning are more likely to become functionally decoupled with the ACC after learning. By contrast, CA1deep exhibited no significant changes across any of these measures further supporting more stable connectivity with ACC (Figure 4D–F). Finally, we examined ACC spike-triggered average of CA1 sublayers. CA1sup showed a significant pre-to-post reduction in baseline-subtracted STA response, whereas CA1deep did not (Figure 1—figure supplement 1). However, the direct comparison of pre-to-post change between CA1sup and CA1deep was not significant (*p = .09*).

**Figure 4.**
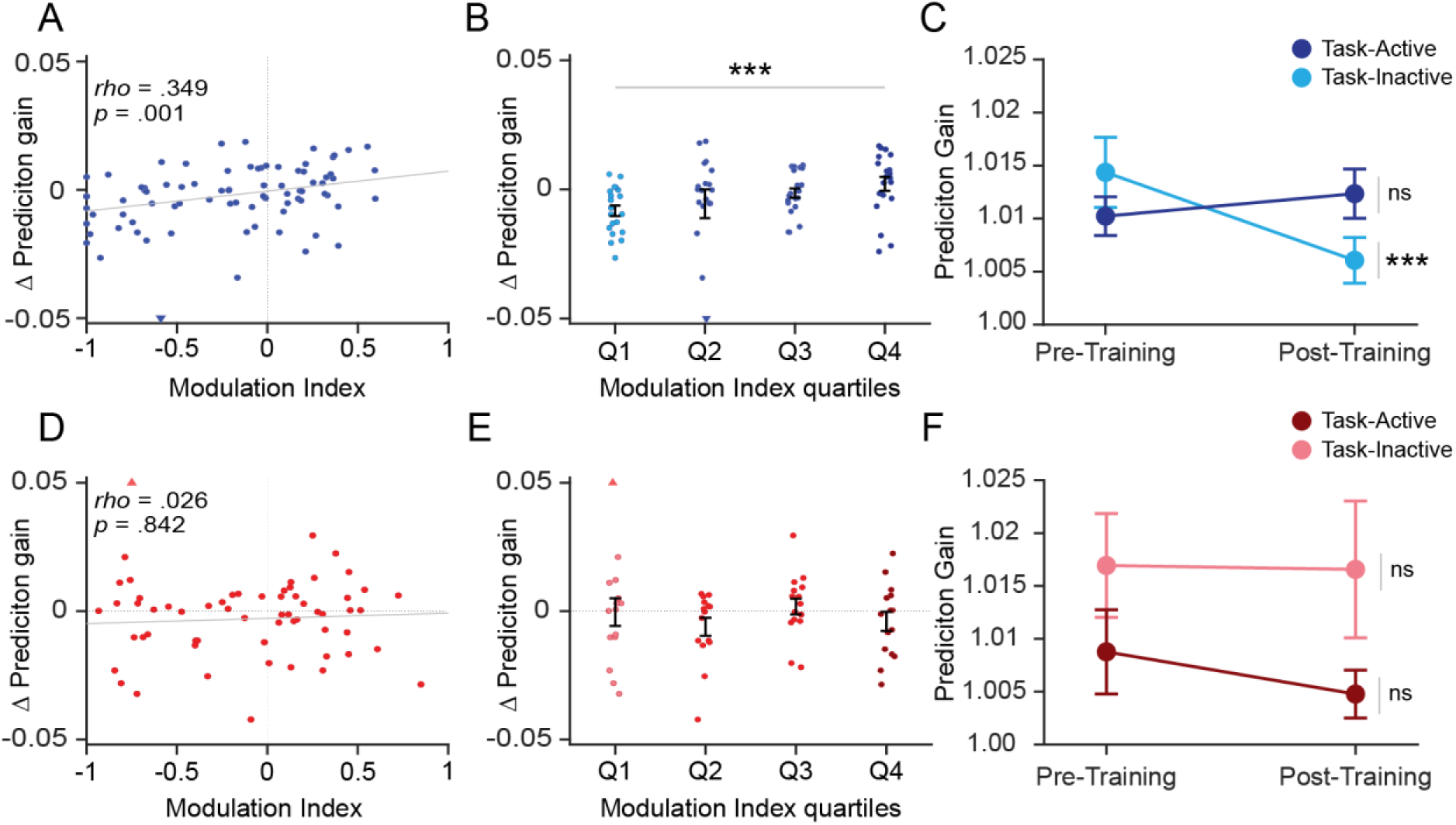
Sublayer differences in ACC→CA1 predictive communication as function of modulation index. **A–C**, PG score differences across CA1sup neurons with respect to modulation index (N = 7 animals; CA1sup: N = 81). **A**, Correlation between CA1sup neurons’ ΔPG and modulation index. Spearman’s Rho revealed a significant positive correlation, *rho* = .349*, p* < .001). **B**, ΔPG as a function of modulation-index quartiles for CA1sup neurons (N = 81). A significant effect of modulation-index quartiles was observed (Kruskal-Wallis, *p* = .014). **C**, PG scores of top and bottom quartiles of CA1sup neurons. A linear mixed-effects model revealed a significant main effect of training session (*p* < .001) and quartile x session (*p* = .002). Post-hoc comparisons revealed a significant pre-to-post decrease in PG score for bottom quartile (Holm corrected, *p* = .002), with no change in top quartile neurons (Holm corrected, *p* = .442). **D–F,** PG score differences across CA1deep neurons with respect to modulation index (N = 6 animals; N = 61 CA1deep neurons). **D**, Correlation between neurons’ PG score and modulation index. Spearman’s Rho revealed a significant positive correlation between ΔPG and modulation index for CA1sup neurons, *rho* = .026*, p* = .842. **E**, ΔPG as a function of modulation-index quartiles for CA1deep neurons (N = 61). CA1deep exhibited no significant effect of modulation-index quartile was observed (Kruskal-Wallis, *p* = .591). **F,** PG scores of top and bottom quartiles of CA1 CA1deep neurons. A linear mixed-effects model did not reveal a significant main effect of training session (*p* = .149) or quartile x session interaction (*p* = .934). Likewise, post-hoc comparisons revealed no changes pre-to-post PG scores for either quartile (Bottom quartile: N = 15, Holm corrected, *p* = .906; Top quartile: N = 15, Holm corrected, *p* = 1). A linear mixed-effects model revealed a trend-level Modulation Index × Sublayer interaction (*p* = .063). Error bars indicate ± SEM.

Overall, ACC→CA1sup predictive relationships were less stable across pre- and post-training sleep, with learning-related reductions occurring specifically among task-inactive CA1 neurons. In contrast, ACC→CA1deep communication remained largely stable across learning. Together, these findings indicate that learning selectively remodels ACC communication with CA1sup neurons, whereas ACC→CA1deep communication is comparatively preserved.

### Optogenetic stimulation of the ACC preferentially inhibits CA1sup neurons

To determine whether the ACC can directly influence the activity of CA1sup neurons, we performed optogenetic stimulation of ACC neurons in a separate cohort of mice while recording CA1 neurons during sleep. We microinjected AAV-CaMKII-ChR2 into the ACC and implanted an optic fiber above the injection site, alongside a recording tetrode array implanted into the CA1 ipsilaterally (Figure 5A—figure supplement 1). After allowing time for viral expression, we administered 4-pulse, 25-Hz optogenetic stimulations during sleep to examine how CA1 neurons responded to ACC stimulations. CA1sup neurons displayed prolonged activity changes, primarily suppression, whereas CA1deep showed limited responses (Figure 5C). To quantify CA1 sublayer differences, we normalized CA1 firing rates across the stimulation window (0–4 sec from stimulation onset) and compared pre-stim to post-stim normalized firing rate (Figure 5D). We found that CA1sup firing rate was significantly lower than CA1deep neurons during the 4 sec post-stim window. Of note, we replicated these stimulation parameters during wakefulness and found more muted responses across each sublayer, suggesting communication between ACC and CA1 is state-dependent (Figure 5—figure supplement 2). Overall, activating ACC produces long-lasting inhibition of CA1sup with limited impact over CA1deep neurons, supporting a selective ACC→CA1sup communication during sleep.

**Figure 5.**
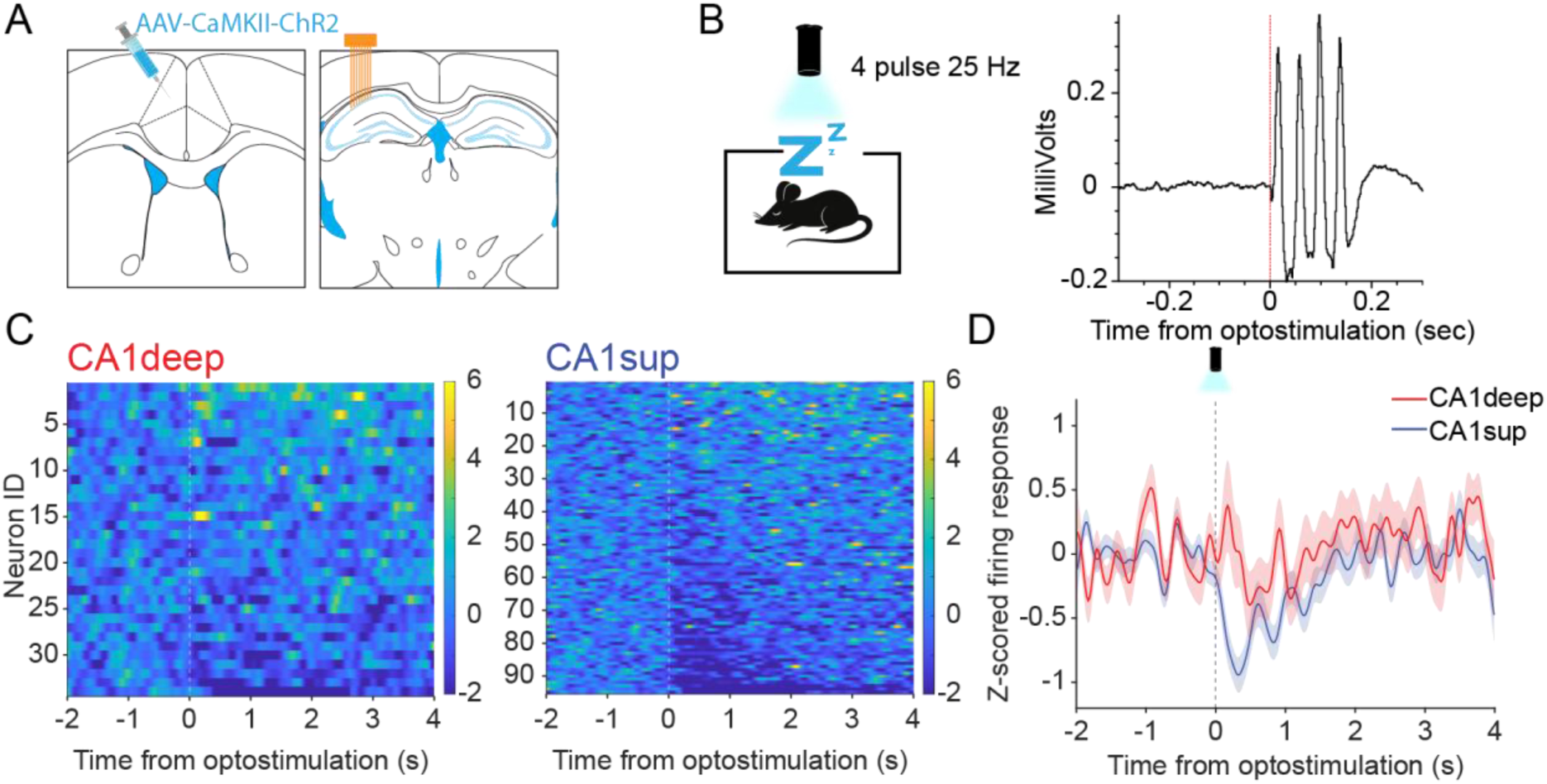
Optogenetic stimulation of the ACC preferentially inhibits CA1sup neurons. **A,** Schematic of AAV injection and microdive implant. **B**, Left, schematic of optogenetic manipulation. Four pulse 25-Hz stimulations were performed during home-cage sleep. Right, representative CA1 LFP response to ACC stimulations. A fast peak, indicating maximal inhibition, occurs approximately 13 ms after stimulation onset. **C**, Heatmap activity CA1deep (N = 34) and CA1sup neurons (N = 96). **D**, Linear mixed-effects model revealed a significant Sublayer × Window interaction for the CA1 activity following ACC stimulation, *p* = .022. There was also an overall sublayer effect, *p* = .037. Post hoc between-sublayer comparisons showed that the strongest difference occurred in the 0–1 sec window, with CA1sup neurons showing greater suppression than CA1deep (*p* < .01, FDR-corrected, *q* = .028). Other time windows were not significantly different after FDR correction. Shaded region indicates mean ± SEM.

### Optogenetic stimulation of the ACC differentially affects CA1 interneurons

Within CA1 lies a complex network of interneurons, which play critical roles in memory processes (Tzilivaki et al., 2023). We aimed to understand how local interneurons within CA1 respond to ACC stimulations. We first investigated the responses of fast-spiking putative parvalbumin (PV) interneurons. Their firing properties have been well characterized and have been shown to display increased activity during ripples (Figure 6 A–C; see Methods for details) (Klausberger & Somogyi, 2008; Opalka et al., 2020; Varga et al., 2014). Upon stimulation of the ACC during sleep, we discovered a robust inhibition of PV interneurons (Figure 6D, E). Like CA1sup neurons, PV suppression was delayed (median latency = 97.5 ms, IQR = 37.5–295 ms) and prolonged, demonstrating that the ACC can drive sustained suppression of both CA1sup and PV interneurons (Figure 5C; Figure 6E; Figure 6—figure supplement 1E).

**Figure 6.**
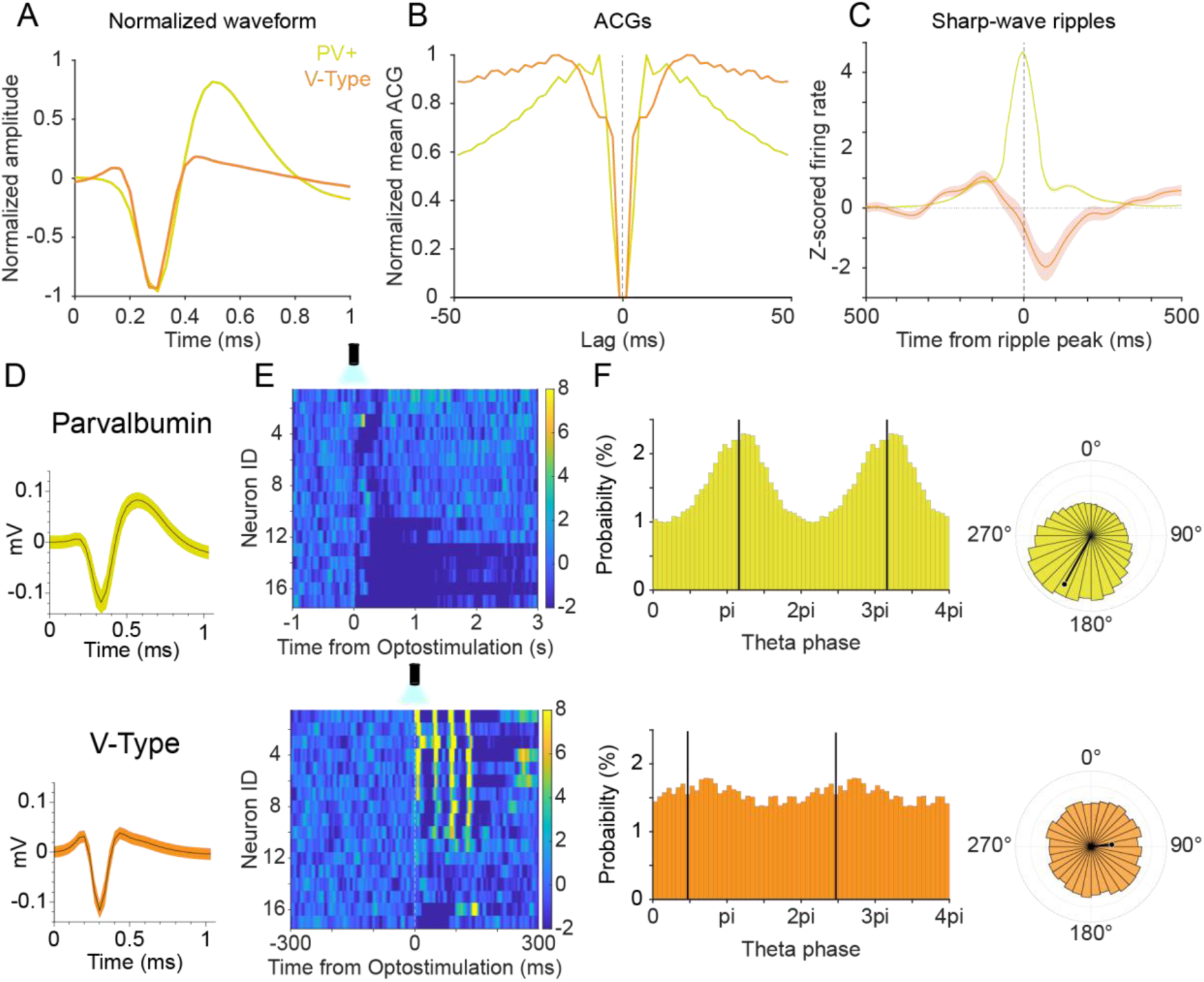
Optogenetic stimulation of the ACC differentially affects CA1 interneurons. **A**, Normalized waveform of putative PV (N = 17) and V-type interneurons (N = 17). **B**, Autocorrelograms (ACGs) of putative PV and V-type interneurons across all spikes. ACGs were calculated for each putative PV and V-type interneuron, then averaged across all neurons within each group. **C**, Normalized firing across population PV and V-type interneurons during sharp-wave ripples. Mann-Whitney U revealed a significant difference in activity during sharp-wave ripples across interneurons, *p* < .001. **D**, Top, representative waveform of putative PV interneuron. Bottom, Representative waveform of a V-type interneuron. **E,** Top, Heatmap of PV interneurons spiking activity following stimulation of the ACC (bin size, 5 ms; N = 17). Bottom, Heatmap of V-type interneurons spiking activity following stimulation of the ACC (bin size, 1 ms; N = 17). **F**, Theta Phase-locking analysis and per-neuron circular statistics for CA1 interneurons during active wakefulness theta. Top, PV interneuron phase-locking response during theta. PV interneurons showed significant phase locking at 28.8° relative to trough, *r* = -.795*, Rayleigh p* < .001. Bottom, V-type interneuron phase-locking response during theta. V-type interneurons showed a per-neuron circular mean preferred phase of 84.5° relative to trough, however, this was not significant, r *= .385, Rayleigh p* = .079. Per-neuron circular statistics showed that PV and V-Type interneurons differed in preferred theta phase (Watson-Williams, *p* < .001) and phase-locking strength (Mann-Whitney U test, *p* < .001). Shaded region indicates mean ± SEM.

Notably, CA1 LFPs exhibited a laser-evoked response peaking roughly 13 ms after ACC stimulation, which precedes many of the responses seen in PV interneurons and CA1sup neurons (median latency = 350 ms, IQR = 130–830 ms) (Figure 5B, C; Figure 6E; Figure 6—figure supplemental 1E). This suggests the presence of a different population of CA1 neurons that respond on a scale similar to that seen in the CA1 LFP. Throughout our recording sessions, we noticed a reoccurring neuron type that showed a V-shape spike waveform and distinct autocorrelegram (Figure 6A, B). Unlike PV interneurons and pyramidal neurons, these V-type interneurons decreased activity during ripples (Figure 6C). Most notable, however, was their robust, low-latency activation to ACC stimulations (median latency = 10.5 ms, IQR = 8.1–13.3 ms) (Figure 6D; Figure 6—figure supplement 1E). Overall, these V-type interneurons had a narrow spike width, were fast spiking (Hz), and displayed a distinct inter-spike interval pattern compared to PV interneurons and pyramidal cells (Figure 6—figure supplement 1). Unlike PV interneurons, V-type interneurons did not exhibit significant theta phase locking, instead firing more uniformly across the theta cycle, and displayed a significantly lower burst index (Figure 6F; Figure 6—figure supplement 1D). Given that V-type interneuron activation precedes suppression of pyramidal cells and PV interneurons after ACC stimulation, it is possible that V-type interneurons mediate the inhibition seen in pyramidal cells and PV interneurons. However, further studies are needed to identify and test whether these neurons influence CA1 network activity.

## Discussion

Balancing network excitability with suppression after learning has long been thought to be essential for memory formation, with sleep serving a key homeostatic role (Liu et al., 2010; Tononi & Cirelli, 2003; Tononi & Cirelli, 2006; Watson et al., 2016) Excessive excitability and overly strengthened synapses can hinder neuronal flexibility and saturate the capacity for learning (Balduzzi & Tononi, 2013; Gava et al., 2024; Norimoto et al., 2018). Additionally, underregulated excitability may pathologically link multiple memory traces, leading to memory overgeneralization (Jinde et al., 2012; Kim et al., 2021). Therefore, regulatory mechanisms that rebalance the excitability of neurons to enable neuronal flexibility are essential for memory formation (Gava et al., 2024; Karaba et al., 2024). CA1sup neurons have been shown to be particularly susceptible to memory rigidity (Gava et al., 2024; Hall & Wang, 2023). While activation of these neurons is necessary for the encoding of new memories, their regulation and timely suppression appear equally important to evoke flexibility and enable *de novo* consolidation within the hippocampal network (Gava et al., 2024; Harvey et al., 2023). Our results reveal a rebalancing of communication between the ACC and CA1 following learning. This was most pronounced for CA1sup neurons as functional connectivity was dynamically reorganized between pre- and post-training sleep. In particular, pre-existing ACC-to-CA1sup communication was weakened for task-inactive neurons after learning. Furthermore, we found that optogenetic ACC stimulations preferentially inhibited CA1sup neurons. Overall, our study uncovers an ACC→CA1sup line of communication that adapts to learning, potentially playing a role in the rebalancing of network excitability for memory formation.

### Learning selectively weakens ACC coupling with task-inactive CA1sup neurons

We implemented GLM machine learning to decode CA1 neuronal firing rates during ripples based off the spiking rates of large ACC neuronal populations preceding ripples (Rothschild et al., 2017). The underlying principles of this analysis posit that if ACC activity preceding ripples influences subsequent activity in CA1 during ripples, the specific activity pattern of ACC neurons (predictor cells) should hold relevancy over the outcome of CA1 neurons (predicted cells). We found this to be true across all sleep sessions and neuron types, with real ACC spiking data showing significantly higher prediction gain compared to shuffled data, indicating an information flow from ACC to CA1 surrounding ripples. Interestingly, prediction gain was weakened following learning specifically for task-inactive CA1sup neurons, suggesting a selective reorganization of communication with CA1sup neurons according to their recruitment during training. However, questions remain over exactly how the ACC influences CA1sup activity and why this information flow rebalances after learning.

One possible explanation is that ACC→CA1sup communication before training may reflect encoding of past experiences. Research has demonstrated that the ACC has a limited role in consolidation immediately after learning (Junyu Liu et al., 2024). Instead, memories become ACC-dependent beginning typically two days after learning (Einarsson & Nader, 2012; Frankland et al., 2004; Junyu Liu et al., 2024). It is possible that heightened pre-existing communication between ACC and CA1sup is reflective of consolidating past experiences. If true, decreased communication between regions may be due to the network’s shift towards prioritizing the encoding/consolidating of the new experience and thus the role of the ACC to CA1, consolidating past memories, is reduced (Diekelmann & Born, 2010; Huelin Gorriz et al., 2023; Wilson & McNaughton, 1994). From a broader perspective, ACC→CA1 could function as part of default mode network activity, and learning induces network shift toward the newly acquired memory (Buckner et al., 2008; Raichle, 2015).

Alternatively, ACC→CA1 communication may contribute to the homeostatic downscaling of memory-unrelated synapses during sleep. Evidence finds that slow-wave sleep is strongly linked to downscaling of non-learning related neuron activity (Gulati et al., 2017; Liu et al., 2010; Tononi & Cirelli, 2003; Tononi & Cirelli, 2006; Watson et al., 2016). Slow-wave sleep ripples in particular depotentiate memory unrelated synapses (Gulati et al., 2017; Norimoto et al., 2018). In our study, we find that the learning-related reduction in communication between ACC and CA1 was selective for task-inactive neurons. Therefore, ACC→CA1sup communication may not simply weaken following learning but rather becomes selectively disengaged from task-inactive neurons, enabling homeostatic downscaling while preserving behaviorally relevant synapses (Liu et al., 2010; Norimoto et al., 2018; Tononi & Cirelli, 2003; Tononi & Cirelli, 2006; Watson et al., 2016). Interestingly, behavioral recruitment alone cannot account for the observed remodeling of ACC→CA1 communication, as CA1sup and CA1deep neurons did not display significant differences in task-related activity (Figure 2—figure supplemental 2). Instead, these learning related changes of task inactive neurons appear sublayer specific.

A potential mechanism supporting this selectivity is suggested by our optogenetic findings, which revealed preferential suppression of CA1sup neurons following ACC stimulations. ACC-mediated inhibition may contribute to the gradual disengagement of subsets of CA1sup neurons overtime as memories transition toward long-term storage (Karaba et al., 2024; Raven & Aton, 2021). Such a mechanism would align with ACC’s known role in systems consolidation and be particularly useful in the regulation of CA1sup neurons as they have been shown to be susceptible to memory rigidity and overgeneralization (Einarsson & Nader, 2012; Frankland et al., 2004; Gava et al., 2024; Junyu Liu et al., 2024). Future experiments casually manipulating this communication are needed to determine whether ACC activity directly influences the downregulation of CA1sup activity during training, or whether these regions are correlated partners whose activity is ultimately mediated by an external brain region.

As research investigating CA1 heterogeneity continues to gain attention, functional distinctions across different behaviors have emerged (Hall & Wang, 2023). Previous research has found that CA1deep neurons are more involved in reward and sensory landmark encoding, while CA1sup neurons are more involved in context and spatial encoding (Danielson et al., 2016; Geiller et al., 2017; Harvey et al., 2023). Moreover, CA1deep place cells are more likely to remap in response to local cue changes than CA1sup place cells (Esparza et al., 2025). There have been conflicting reports over which sublayer is more involved in learning and replay and which neurons are more plastic or rigid in response to learning (Berndt et al., 2023; Geiller et al., 2017; Grosmark & Buzsáki, 2016; Hall & Wang, 2023; Harvey et al., 2023). We speculate that it may depend on the behavioral task and the relevant upstream circuits that differentially recruit CA1 sublayers (Masurkar et al., 2017). Past reports, which have shown that CA1sup neurons are more involved in memory replay, implemented spatial learning tasks (Berndt et al., 2023; Harvey et al., 2023). In contrast, CA1deep neurons have been shown to be more important for encoding non-spatial aspects of memories, such as reward (Danielson et al., 2016; Harvey et al., 2023). In our study, we did not observe clear differences in sublayer responses to contextual fear conditioning. Instead, we uncovered a pathway specific difference with ACC→CA1sup displaying a more plastic-like phenotype in response to learning compared to ACC→CA1deep. These findings, along with past reports, suggest that functional differences between CA1 sublayers may be better understood in the context of the specific behavioral and upstream circuits engaged during learning rather than as fixed “plastic” or “rigid” properties of the sublayers themselves (Masurkar et al., 2017). In fact, recent findings further complicate this notion, demonstrating sublayer differences emerging between two classes of ripples that differ based on current source density profiles (Castelli et al., 2025). Ultimately, future studies are needed to fully dissect the functional differences among CA1 sublayers and their inputs during ripples and across behaviors.

Lastly, our CA1 sublayer classification does come with its own caveats. Tetrode identification of CA1 sublayers is not a trivial matter. We implemented guidelines informed by past research (see Methods) to help ensure we isolated CA1sup and CA1deep groups, only including neurons where we had the greatest confidence (Berndt et al., 2023; Mizuseki et al., 2011). In doing so, we excluded neurons which classification was uncertain. It is possible this excludes some meaningful populations of CA1 sublayers. Additionally, despite our approach, some cross-inclusion of sublayers may remain. Therefore, interpretations of CA1 sublayers difference should be considered with these limitations in mind. That said, the number of neurons and animals tested does provide overall confidence regarding our results. Future studies investigating this ACC to CA1 sublayer specific line of communication would benefit from the use of silicon probes or neuropixels that enable precise radial localization. Another caveat to mention is that both the ACC and CA1 are involved in the stress response (Kim et al., 2015; Lamotte et al., 2021). Future experiments utilizing appetitive learning paradigms, rather than the aversive contextual fear conditioning used here, will help disentangle learning-related remodeling of ACC→CA1 communication from changes driven by stress.

### Optogenetic stimulation of the ACC inhibits CA1sup neurons and PV interneurons

We found that direct stimulation of ACC excitatory neurons inhibited CA1sup and PV interneurons. Notably, these responses were delayed for CA1sup and PV interneurons, indicating polysynaptic connectivity (Piñol et al., 2012). Additionally, both neuron types exhibited sustained suppression in response to stimulations suggesting that the ACC may have an indirect and broad influence over CA1 network activity rather than a direct and transient role. Sustained inhibition over certain populations of pyramidal neurons and PV interneurons may support a rebalancing of network excitability, especially during sleep (Diekelmann & Born, 2010; Norimoto et al., 2018; Tononi & Cirelli, 2003).

The identification of V-type neurons may help explain responses seen in CA1sup and PV interneurons. These neurons displayed the lowest latency responses to stimulations and produced brief excitatory bursts indicative of an initial response to ACC stimulations (Piñol et al., 2012). These interneurons decreased their activity during ripples, a pattern that overlaps with *Sncg-*expressing CCK basket cells and inhibitor of DNA binding 2 (ID2) neurons (Dudok et al., 2021; Karaba et al., 2024; Valero et al., 2025; Vancura et al., 2023). It is possible that V-type neurons may overlap with these interneurons. Correspondence with CCK cell activity is of particular interest, given the role of CCK interneurons in CA1 circuitry. Namely, CCK interneurons are known to target CA1sup and PV interneurons and induce long-term excitability changes as seen in our optogenetic experiments (Hefft & Jonas, 2005; Karson et al., 2009; Klausberger et al., 2005; Valero et al., 2015). More recently, CCK interneurons have been identified as playing a causal role in regulating the excitability of CA1 neurons during ripples (Karaba et al., 2024). Together, this suggests a possible ACC influence on CA1 CCK interneurons, which provides sustained inhibition of CA1 pyramidal and PV interneurons during sleep in facilitating memory formation. One discrepancy, however, is that both CCK and ID2 neurons exhibit theta phase preference, which we found no evidence for in V-type interneuron (Klausberger et al., 2005; Valero et al., 2025). Ultimately, our interpretations of V-type identity remain speculative as further studies are needed to characterize their activity, and casual experiments are required to completely elucidate their role.

Another important caveat to mention is that it remains an ongoing debate whether ACC directly projects to CA1 (Andrianova et al., 2023; Rajasethupathy et al., 2015; Shi et al., 2022). One lab reported monosynaptic ACC to CA1 connection (Rajasethupathy et al., 2015), while another lab replicated those same experiments but were unable to come to the same conclusions (Andrianova et al., 2023). Further studies report no direct connection (Shi et al., 2022). The contention in connectivity may arise from differences in targeting strategies, injection coordinates, or viruses used. Our findings reported an excitatory response in CA1 V-type interneurons in response to ACC stimulations, proposing another possibility for ACC→CA1 connectivity. Interestingly, V-type interneurons responded with low-latency as fast as 4.2 ms after stimulations, compatible with monosynaptic timing (Cho et al., 2013; Petreanu et al., 2007; Wang et al., 2009). If the ACC→V-Type connection was a monosynaptic pathway, it could help explain discrepancies in the field, as the relative sparsity of V-type interneurons may reduce the likelihood of detecting ACC→CA1 connectivity. However, future anatomical studies are needed to conclusively determine connectivity.

Alternatively, ACC→CA1 communication may be mediated by multiple intermediate structures (Behzadi et al., 1990; Oh et al., 2014; Shi et al., 2022; Souza et al., 2022). The ACC sends monosynaptic projections to the nucleus reuniens (RE) and median raphe (MnR), both of which project directly to CA1 (Oh et al., 2014; Shi et al., 2022). The RE has a known role in contextual discrimination learning and memory specificity (Ramanathan & Maren, 2019; Ramanathan et al., 2018; Ratigan et al., 2023; Silva et al., 2021; Xu & Südhof, 2013). Interestingly, RE→CA1 activity tuned to immobility (freezing) emerges only after shocks are presented, suggesting a learning induced modification between regions, similar to that seen in our ACC–CA1 data (Ratigan et al., 2023). As for MnR, it receives dense inputs from the ACC (Behzadi et al., 1990; Souza et al., 2022), and its projections to the CA1 are predominantly glutamatergic (Jackson et al., 2009; Senft et al., 2021; Szonyi et al., 2016). Notably, these glutamatergic MnR inputs directly target CA1 interneurons, including CCK basket cells (Miettinen & Freund, 1992; Morales & Bloom, 1997; Senft et al., 2021), while avoiding PV interneurons and pyramidal neurons (Acsady et al., 1993; Freund et al., 1990; Halasy et al., 1992; Miettinen & Freund, 1992; Papp et al., 1999; Turi et al., 2019). This connectivity suggests that the ACC may indirectly modulate CA1 activity through the MnR, potentially suppressing PV interneuron and pyramidal neuron activity via local inhibitory circuits, thereby contributing to the regulation of hippocampal oscillations and memory consolidation (Huang et al., 2022; Wang et al., 2015). Ultimately, future experiments combining pathway-specific manipulations with simultaneous recordings will be necessary to distinguish direct from polysynaptic mechanisms.

## Methods

### Mice

Male C57BL/6 mice were purchased from the Jackson Laboratory (stock #000664). Mice were 3–4 months old at the time of surgery; after surgery, they were singly housed in cages (40 × 20 × 25 cm) containing corn cob and cotton material and kept on a 12 h light/dark cycle with *ad libitum* access to food and water. Experimental procedures were approved by the Institutional Animal Care and Use Committees of Drexel University (protocol # LA-23-740) and were in accordance with the National Research Council *Guide for the Care and Use of Laboratory Animals*.

### Stereotaxic surgery

Surgery procedures were similar to that used in our lab (Huang et al., 2026; Jun Liu et al., 2024; Liu et al., 2021). In brief, mice were anesthetized with ketamine/xylazine mixture (∼100/10 mg/kg, i.p.) and kept on a heating pad at 37°C. For *in vivo* electrophysiology recording, mice received implantation of two electrode arrays (8 tetrodes each) into the ACC and CA1, respectively (Lin et al., 2006). For optogenetic stimulation, mice received intra-ACC microinjection of AAV viruses (AAV1-CKIIa-ChR2-GFP; 0.250 μl; ∼10^13^ GC/ml; *Addgene* 105669) and implantation of one optic fiber (diameter 200 μm) slightly above the injection site; meanwhile, they received implantation of 8 tetrodes into the CA1 unilaterally on the ipsilateral side. AAV vectors were microinjected through a syringe pump (*World Precision Instruments*) over 5 min (50 nL/min), with an additional 5 min waiting period before removal of the injection needle (34 gauge, beveled). ACC coordinates from Bregma were AP 0.9 mm, ML 0.3 mm, DV 1.0 mm; CA1 coordinates from Bregma were AP –2.1 mm, ML 1.7 mm, and DV 1.1 mm.

### *In vivo* electrophysiology

Each tetrode consisted of four wires (90% platinum 10% iridium; 18 μm diameter; *California Fine Wire*). A microdrive was used to couple with the electrode array, similar to that used in our lab (Jun Liu et al., 2024; Liu et al., 2021; Wang et al., 2015). Neural signal was preamplified, digitized, and recorded using a *Blackrock Neurotech* CerePlex. The local field potentials (LFPs) were digitized at 2 kHz and filtered at 500 Hz low cut; spikes were digitized at 30 kHz and filtered between 600–6000 Hz. The tetrode arrays were gradually lowered daily until we recorded clear ripples and a substantial number of neurons; otherwise, mice were excluded from the study. The recorded spikes were sorted manually using *Plexon* Offline Sorter. For dual-site experiments, spikes from 10 mice were used for analyses in this study; the neuron numbers in ACC and CA1 were 48/27, 35/50, 28/12, 32/27*, 22/29, 26/17, 28/33,19/17, 27/47, and 38/32 respectively. For optogenetic and *in* vivo electrophysiology experiments, 5 mice were used. The neurons numbers in CA1 were: 35, 30, 10, 38, and 24.

* This animal is missing the recording file during training and therefore was excluded from analyses comparing task active/inactive and firing rate ratios during training.

### Optogenetics manipulations

Following surgery, mice were given 1 week for recovery and viral expression development. During recording, mice received 4-pulse 25-Hz stimulations (20-sec inter-trial interval). Laser stimulations ranged between 5–8 mW (473 nm wavelength). Laser intensity was tuned for each recording session until stimulation evoked a >0.2 mV change in CA1 LFP.

### Ripples

Local field potentials were band-pass filtered between 100–250 Hz and ripple envelope was smoothed with a Gaussian kernel (S.D. = 4 ms) (Karlsson & Frank, 2009). We defined ripple amplitudes as the peak values of ripple envelopes. Only ripples with amplitudes exceeding 5 s.d. above the mean were used for analyses. For each ripple, onset and offset were taken as the earliest and latest time points where the ripple amplitude exceeded 1 SD above the mean around the ripple peak. Ripple duration was the difference between ripple onset and offset. Ripples less than 20 ms were excluded from the analysis.

### Modulation and Burst index

Modulation index was a firing rate ratio calculated by comparing spiking during training with pre-training sleep baseline) (Girardeau et al., 2017; Karaba et al., 2024; Oliva et al., 2020). Equation: Modulation index = (Train – Pretrain) / (Train + Pretrain), in which Train and Pretrain are the average spikes per second for each CA1 neuron calculated during training and pre-training sleep, respectively. Burst index was defined as the fraction of consecutive inter-spike intervals shorter than 6 ms across the full session recording (Harris et al., 2001).

### GLM decoding

We constructed generalized linear models (GLMs) with a log link function to predict spike counts of individual CA1 neurons during ripples based on population spike counts in ACC across specific time windows (Jun Liu et al., 2024; Rothschild et al., 2017). Spike counts of each ACC neuron were binned in 200-ms bins relative to ripple onset: −600 to −400 ms, −400 to −200 ms, −200 to 0 ms to predict CA1 activity 0–100 ms. We randomly partitioned the ripples into two approximately two equally sized datasets: one of them was used to train the GLM decoder, and the other was used for testing. The model derived from the training phase was applied to the ACC population spike data in the test set, yielding predictions for the predicted CA1 spike counts across ripples. We calculated a prediction error for each CA1 neuron that was the mean absolute error between the predicted spike rate and real spike rate. We replicated this same analysis for shuffled data; wherein shuffled error distribution was generated by randomly permuting the predicted CA1 spike counts across ripple events. Prediction gain scores were derived by dividing shuffled prediction error by the real prediction error. Slow-wave sleep was determined by animals’ immobility and the theta (6–12 Hz) /delta (1–4 Hz) ratio extracted from the local field potentials. A ratio of 1 or lower was identified as SWS, only epochs greater than 10 min were selected (Wang et al., 2015)..

### ACC activity surrounding ripples

To calculate significant differences in ACC activity surrounding ripples, we computed peri-ripple event histograms (smoothed with a three-bin Gaussian filter; bin size, 5 ms) for individual ACC neurons. We then compared ACC spiking activity between two 250 ms windows starting either –200 ms or –2000 ms (baseline) before ripple onset. 235 neurons (94%) showed significant increase from baseline (Wilcoxon-signed rank *p* < .01). Pre- or post-ripple peaks were determined whether a neurons firing rate peaked before (–200 to 0 ms) or after (0–200 ms) ripple onset.

### Classification of CA1 pyramidal and interneurons

Putative excitatory (pyramidal) neurons and parvalbumin fast-spiking interneurons were classified based on spike waveform, firing rate, and spike width (Opalka et al., 2020) (Figure 6—figure supplement 1A–C). V-type interneurons were classified based on waveform, firing rate, spike width, and decreased ripple-correlated activity (Figure 6—figure supplement 1A–C). For sublayer classification, we examined the mean amplitude of the sharp wave deflection at maximum ripple power for each tetrode. Sharp-wave deflections that were greater than 50 μV were classified as CA1deep sublayer, whereas deflections less than –50 μV were classified as CA1sup sublayer. Any deflections between –50 and 50 μV were treated as intermediate and were excluded from sublayer analyses (Figure 3A).

### Response latency to ACC optogenetic stimulations

Neural activity following stimulation onset was binned (CA1sup, 20 ms; PV, 5 ms; V-type, 0.1 ms) and z scored. For PV interneurons and CA1sup neurons, response latency was obtained by selecting the first bin where a neuron showed a difference from the baseline by a z-score of ± 2 or more (minimum for 3 consecutive bins). Because V-type interneurons exhibited substantially shorter response latencies, a higher temporal resolution analysis was performed. Spikes were aligned to four-pulse optogenetic stimulation trains and binned at 0.1-ms resolution. Firing rates during inter-pulse windows were z-scored relative to baseline, and response latency was defined as the time from stimulation onset to the peak response in the window with the strongest activation. Only neurons with a maximal response of z ≥ 5 were included in the latency analysis.

### Spike-theta phase locking

Theta oscillations were obtained from CA1 LFPs filtered at 6–12 Hz using a zero-phase filter in MATLAB (bandpass IIR filter; filter order = 8). Theta phase was estimated by Hilbert transformation of the filtered signal. For each neuron, spike times occurring during selected theta epochs were matched to the nearest LFP sample, and the corresponding instantaneous theta phase was extracted for circular statistical analysis. Phase locking was quantified using CircStat functions (Berens, 2009). We used standard Rayleigh test to determine uniformity of phase distribution (Totah et al., 2013), in which the p value for circular uniformity is calculated as follows:

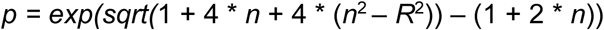

and Rayleigh’s z to determine the strength of spike–LFP phase locking as follows:

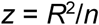

where *R* is the sum of resultant vectors across *n* spikes. Log-transformed Rayleigh z values were calculated log z values and stored for each neuron/group.

### Spike Triggered Average (STA)

For each CA1 neuron, all ACC spikes recorded during slow-wave sleep in that same session were pooled and used as reference events. The spike times of each CA1 neuron were aligned to the pooled ACC spike. Relative CA1 spike times were counted in 10-ms bins over a window spanning −200 to 300 ms relative to the pool ACC spike. Spike counts within each lag bin were summed across all ACC spikes and converted to firing rate (Hz) by dividing by the total number of ACC spikes and the bin width. To normalize each neuron’s lag-resolved STA, the mean firing rate during the −200 to −100 ms pre-ACC-spike baseline window was subtracted from every lag bin for that neuron. Thus, the normalized STA curve estimates the change in CA1 firing rate relative to the pre-ACC-spike baseline as a function of time from ACC spiking. Full-lag significance was assessed using an animal-level sign-flip cluster permutation test (Maris & Oostenveld, 2007). Post-minus-Pre STA difference curves were calculated for each animal, contiguous lag bins exceeding an uncorrected paired-test threshold were grouped into clusters and observed cluster masses were compared against a 5000-permutation sign-flip null distribution.

### Fear conditioning (*in vivo* recording)

The fear-conditioning chamber used in the experiment was a square chamber measuring 25 × 25 × 32 cm, with a 36-bar shock grid floor (*Med Associates*). The behaviors of the mice were recorded using *Blackrock Neurotech NeuroMotive* video system. During training, the mice were first allowed to explore the footshock chamber for 3 minutes. They then received 3 mild footshocks (0.75 mA, 0.5 sec), with a 2-min interval between shocks. About 30 seconds after the last shock, the mice were returned to their home cages. After ∼2 hours of post-training sleep, the mice were placed back in the footshock chamber for a 5-min contextual fear test. Neural activity was recorded continuously, including the pre-training sleep (2–3 hours), training (7.5 min), post-training sleep (2–3 hours), and contextual-fear test (5 min). Freezing was defined as the absence of visible movement expect for respiration as previously described and was scored continuously during the recall test (Anagnostaras et al., 2003; DeLorey et al., 1998). Percent Freezing was defined as time spent freezing divided by total time in recall (or pre-shock baseline during CFC). Additionally, the day prior to fear conditioning, mice were placed into a novel box for 7.5 min; neural activity was recorded across the pre-exposure sleep, exposure, and post-exposure sleep phases, to act as a behavioral control, however, these datasets were not analyzed for this manuscript.

### Histology

To mark the final recording sites, we made electrical lesions by passing 20-second, 10-μA currents through two or more tetrodes. Mice were deeply anesthetized and intracardially perfused with ice-cold PBS or saline, followed by 10% formalin. The brains were removed and postfixed in formalin for at least 24 hours. The brains were sliced into coronal sections of 50-μm thickness using *Leica* vibratome. Sections were mounted with Mowiol mounting medium mixed with DAPI for microscopic examination of electrode placements, viral vector expression, and/or optical fiber placements.

### Statistics

Sample sizes were based on previous similar studies in our labs (Jun Liu et al., 2024; Liu et al., 2021; Wang et al., 2015). Other statistical analyses include Freidman’s test, Wilcoxon signed-rank test, Mann-Whitney U test, Kruskal-Wallis test, Welch’s t-test, independent sample’s t-test, and Student’s *t* test. All statistical tests were two-sided. P-values of 0.05 or lower were considered significant. Non-parametric tests were utilized when normality assumptions failed.

### Data availability

Neural datasets, including all neural spiking and event timestamps, neuron waveforms, and local field potential recordings will be made publicly available on a public repository.

### Code availability

Key customized MATLAB code used in the analysis will be available upon publication of this manuscript.

### Author Contributions

A.F.H. D.V.W. conceptualized the study. A.F.H performed all experiments. A.F.H and D.V.W. wrote the manuscript.

### Financial Disclosures

The authors declare no financial interest or potential conflicts of interest

## Acknowledgments

We thank the members of the Drexel ULAR staff for support and Dr. Wen-Jun Gao, Shayna Singh, Ashley Opalka, and Wenqiang Huang for their thoughtful comments.

## Supplementary Figures

**Figure 1—figure supplement 1.**
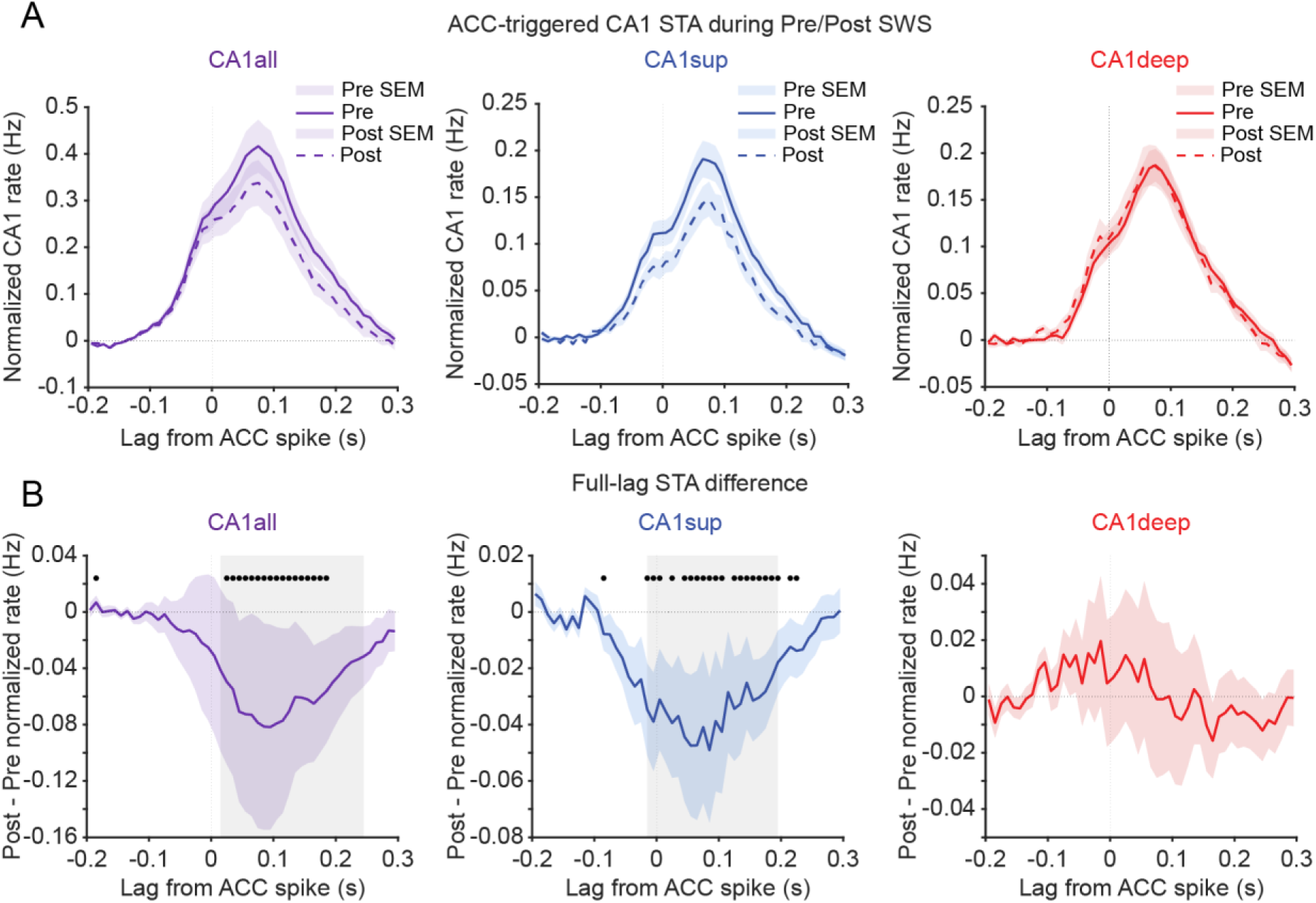
ACC→CA1 coupling decreases for CA1sup neurons, **A**, ACC-triggered CA1 spike-triggered averages during pre- and post-training SWS, normalized to each neuron’s −200 to −100 ms pre-ACC-spike baseline for all CA1 neurons (left; N = 291), CA1sup (middle; N = 89) and CA1deep (right; N = 62). **B**, post-training minus pre-training STA difference curves. Black dots indicate lag bins surviving Benjamini–Hochberg false discovery rate (FDR) correction, and gray shading denotes significant clusters identified by an animal-level sign-flip cluster permutation test. Left, CA1 neurons exhibited a significant negative cluster spanning 15–245 ms after ACC spikes (cluster permutation, *p* = .003). Middle, CA1sup neurons similarly exhibited a significant negative cluster spanning −15 to 195 ms relative to ACC spikes (cluster permutation, *p* = .008). Right, no significant cluster was detected in CA1deep neurons (cluster permutation, *p* = .240). Consistent with the full-lag analysis, animal-level baseline-subtracted STA responses were significantly reduced following learning in CA1 neurons (N = 10, paired *t*-test, *p* = .008) and CA1sup neurons (N = 8, paired *t*-test, *p* = .015), but not in CA1deep neurons (N = 7, paired *t*-test, *p* = .838). Direct comparison of animal-level pre-to-post STA changes between CA1sup and CA1deep was not significant (Mann–Whitney *U* test, *p* = .093). Data not shown.

**Figure 1—figure supplement 2.**
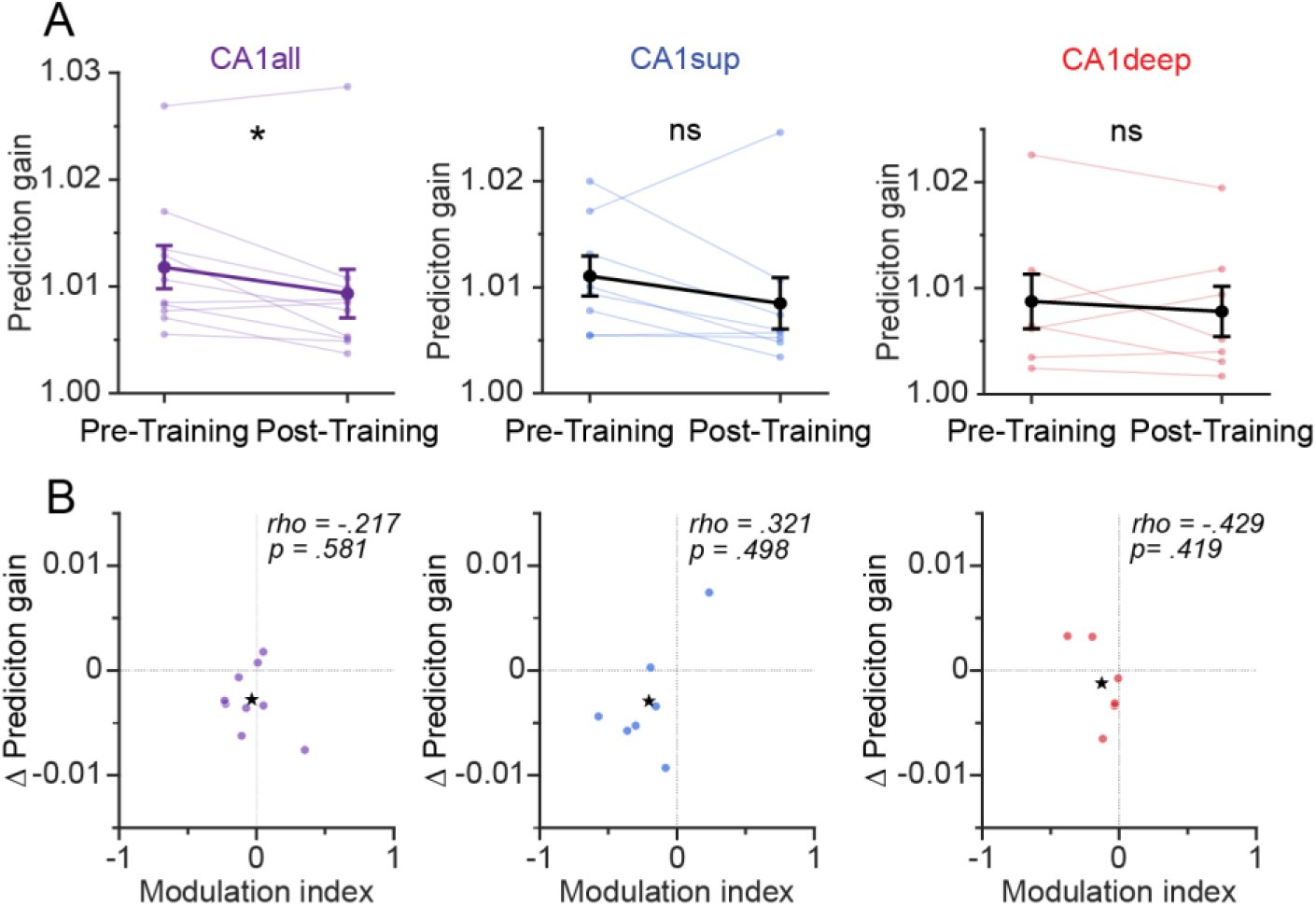
Prediction gain differences per animal, **A**, Prediction gain differences between CA1 neurons across neurons grouped by animal. Left, Prediction gain difference across all recorded CA1 neurons averaged across animals (N = 10) across pre- and post-training sleep. Wilcoxon signed-ranked test revealed a significant pre-to-post decrease in prediction gain across animal, *p* = .048. Middle, prediction gain averaged across all recorded CA1sup neurons per animal (N = 8) across pre- and post-training sleep. Wilcoxon signed-ranked test found no significant differences pre to post decrease in prediction gain across animal, *p* = .250. Right, prediction gain difference across all recorded CA1deep neurons averaged per animal (N = 7) across pre- and post-training sleep. Wilcoxon signed-ranked test found no significant differences in prediction gain for CA1deep neurons across training sessions averaged across animal, *p* = .575. **B,** Correlations between modulation index and ΔPG for all CA1 neurons (N = 9), CA1sup (N = 7), and CA1deep (N = 6) per animal. Left, there was no significant correlation between modulation index ΔPG on a per animal basis for all CA1 neurons (Spearman’s *rho* = -.217*, p* = .581. Middle, there was no significant correlation between modulation index ΔPG on a per animal basis for CA1sup neurons (Spearman’s *rho* = .321*, p* = .498. Right, Likewise, there was no significant correlation between modulation index ΔPG on a per animal basis for CA1deep neurons (Spearman’s *rho* = -.429*, p* = .419). Error bars indicate SEM.

**Figure 2—figure supplement 1.**
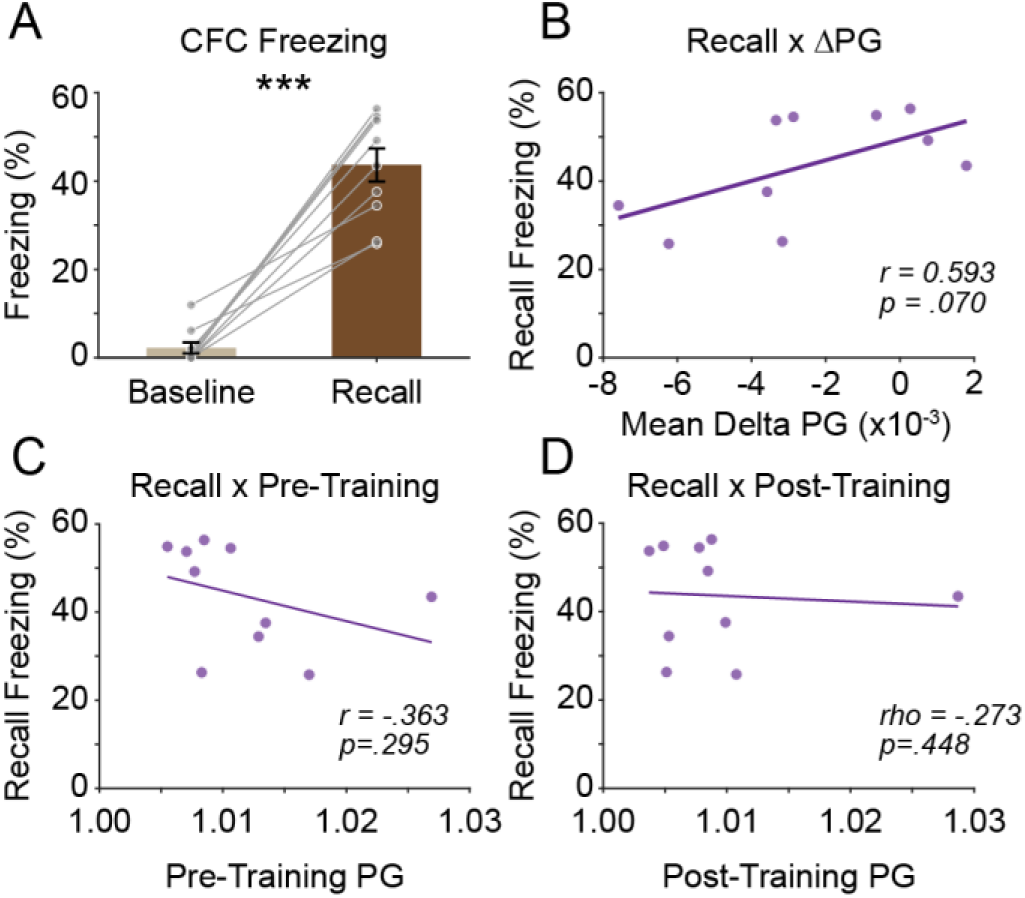
CFC freezing response correlated with Prediction gain, **A**, Time spent freezing during CFC baseline and recall. Paired t-test revealed a significant difference in time spent freezing between BL and recall, *p* < .001 (N = 10 animals). **B**, Correlational analysis between time spent freezing during recall and prediction gain change delta. Pearson’s correlation was not significant between variables *r* = .593*, p* = .070. **C**, Correlation analysis between time spent freezing during recall and pre-training prediction gain (PG) scores. There was not a significant correlation between pre-training PG and freezing during recall, *r* = -.368*, p =* .295. **D**, Correlational analysis between time spent freezing during recall and post-training PG scores. Spearman’s rho did not reveal a significant correlation between post-training PG scores and freezing during recall, *rho* = -.273*, p* = .488. Error bars indicate SEM.

**Figure 2—figure supplement 2.**
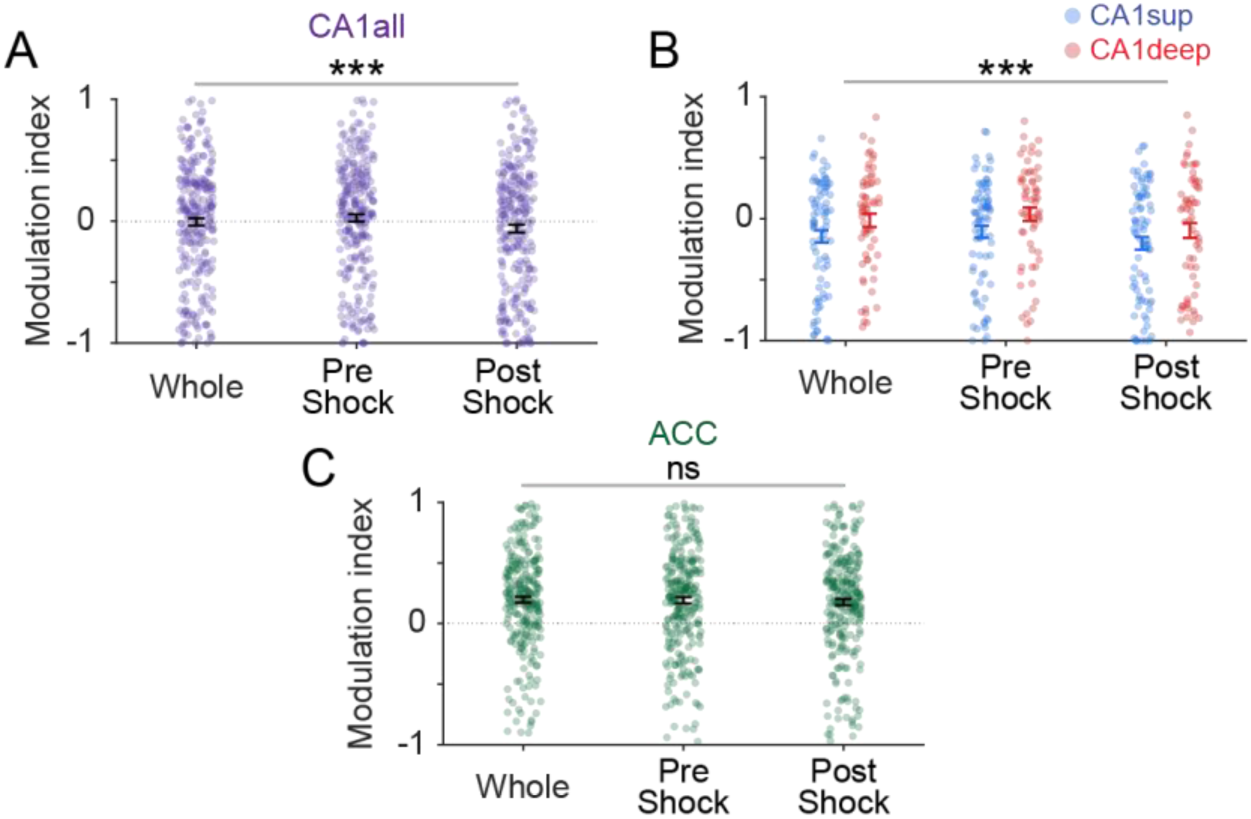
Modulation index changes across CFC and modulation index differences between sublayers. **A–C**, Modulation indexes separated into discrete time windows: whole CFC recording, pre-shock baseline or post-shock windows. **A,** Modulation index for all CA1 neurons across three CFC windows (N = 264). A linear mixed-effects model revealed a significant main effect of window, *p* < .001. Post-hoc paired comparisons showed that modulation index was significantly higher during the pre-shock baseline than whole CFC, (Wilcoxon signed-rank, *p* < .001). Additionally, post-hoc comparisons significantly lower modulation index during the post-shock onset window than both whole CFC and pre-shock baseline, (Both, Wilcoxon signed-rank, *p* < .001). **B,** Modulation index for ACC neurons across three CFC windows (N = 271). A linear mixed-effects model revealed no significant main effect of window, *p* = .140, **C,** Modulation index for CA1 firing activity across three CFC windows for CA1 sublayers (CA1sup: N = 81; CA1deep: N = 61). Linear mixed effect ANOVA model revealed a significant main effect of window for CA1 sublayers, *p* < .001. In CA1sup, modulation index was significantly higher during pre-shock baseline than whole CFC and significantly lower during post-shock onset than both whole CFC and pre-shock baseline, all Wilcoxon signed-rank *p* < .001. In CA1deep, modulation index was also higher during pre-shock baseline than whole CFC, paired t-test *p* = .009, and lower during post-shock onset than both whole CFC, Wilcoxon signed-rank *p* = .003, and pre-shock baseline, Wilcoxon signed-rank *p* = .008. However, there was no significant main effect of sublayer, *p =* .090. Furthermore, Window × Sublayer interaction was also not significant, *p* = .487. Error bars indicate ± SEM.

**Figure 5—figure supplement 1.**
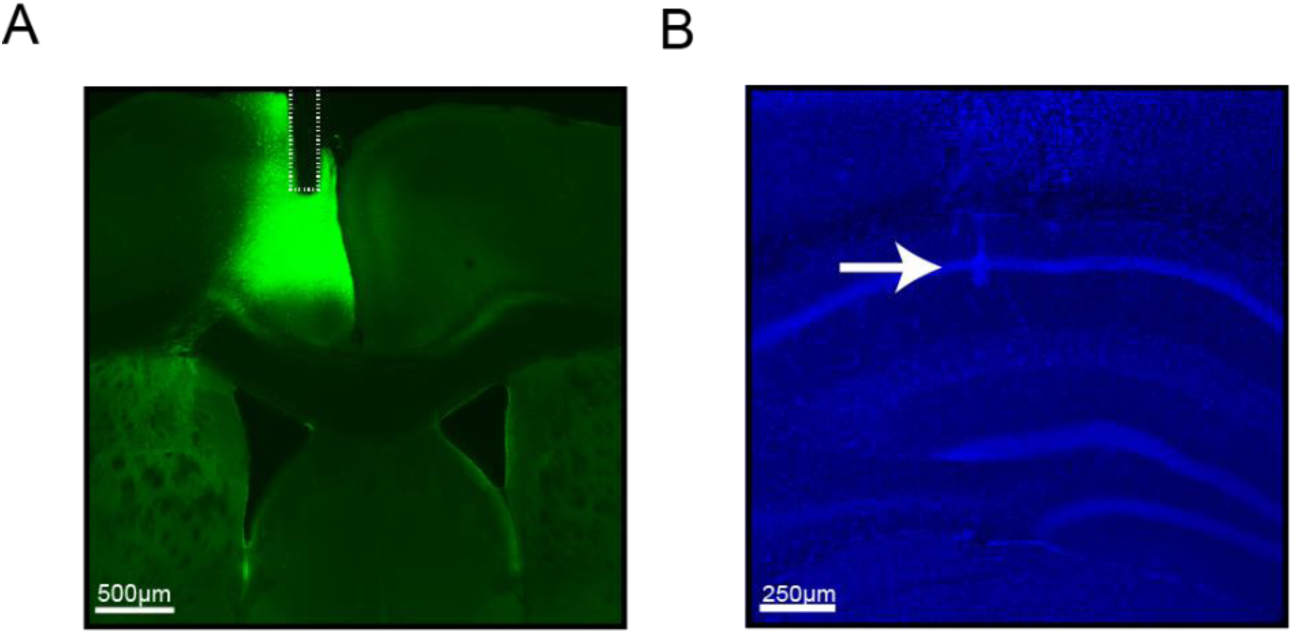
Representative histology for optogenetic experiments. **A**, Optic fiber placement and ChR2 expression. **B,** Tetrode placement for CA1, arrow indicates location of tetrode tip.

**Figure 5—figure supplement 2.**
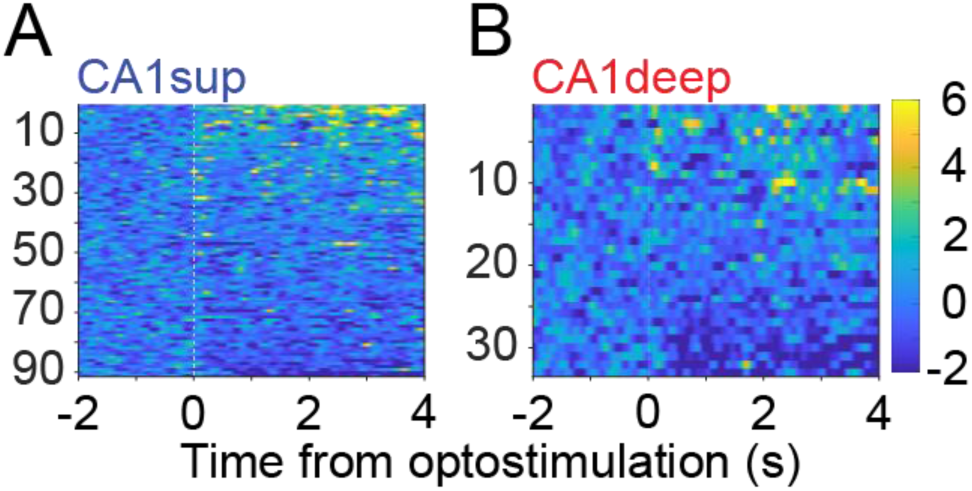
CA1 neuron response to ACC stimulation, **A**, Heatmap for CA1sup neuron spiking (bin 20 ms) during wakefulness ACC stimulations (N = 94). **B,** Heatmap for CA1deepp neuron spiking (bin 20 ms) during wakefulness ACC stimulations (N = 34). Linear mixed effect model revealed no difference between CA1sublayers following ACC stimulation during wakefulness, *p* = .549.

**Figure 5—figure supplement 3.**
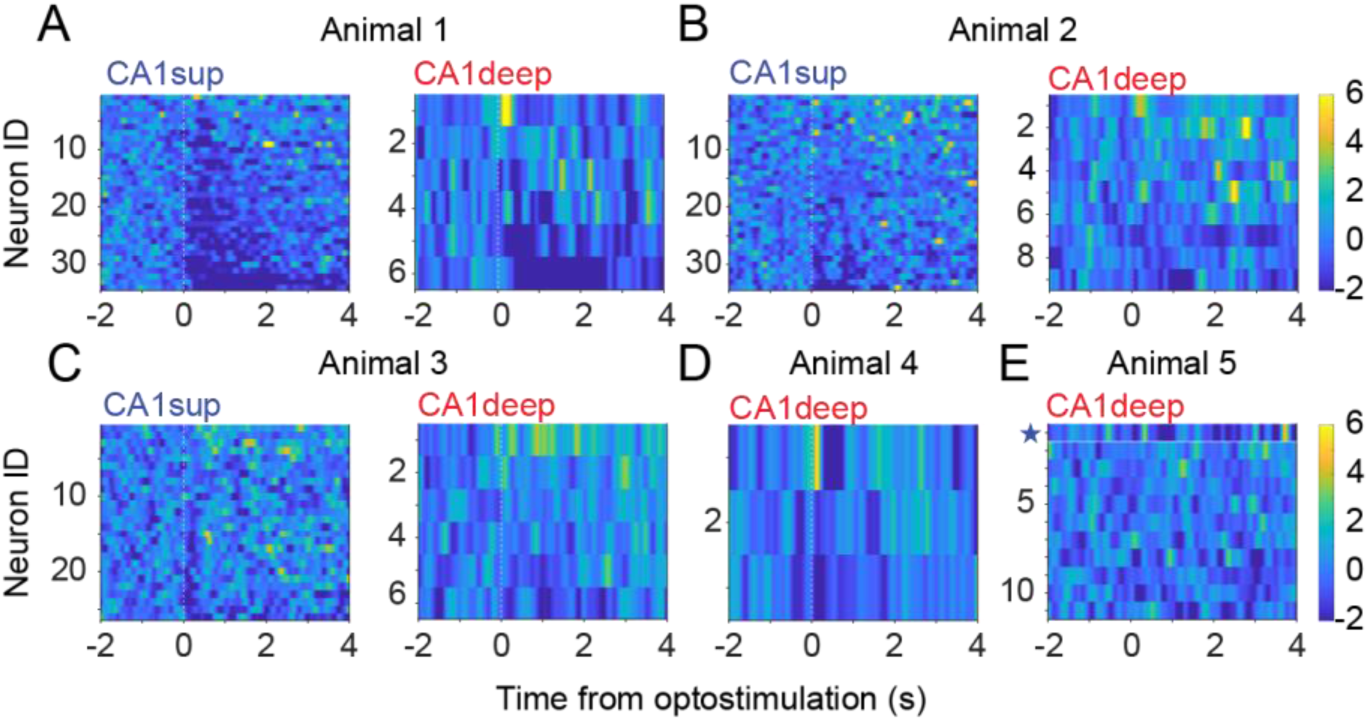
**A–E**, Heatmaps for CA1 pyramidal neuron spiking (bin, 20 ms) separated per animal. **E,** Note, Animal 5 only had one recorded CA1sup neuron. That neuron was added to the CA1deep heatmap (Indicated by blue star).

**Figure 6—figure supplement 1.**
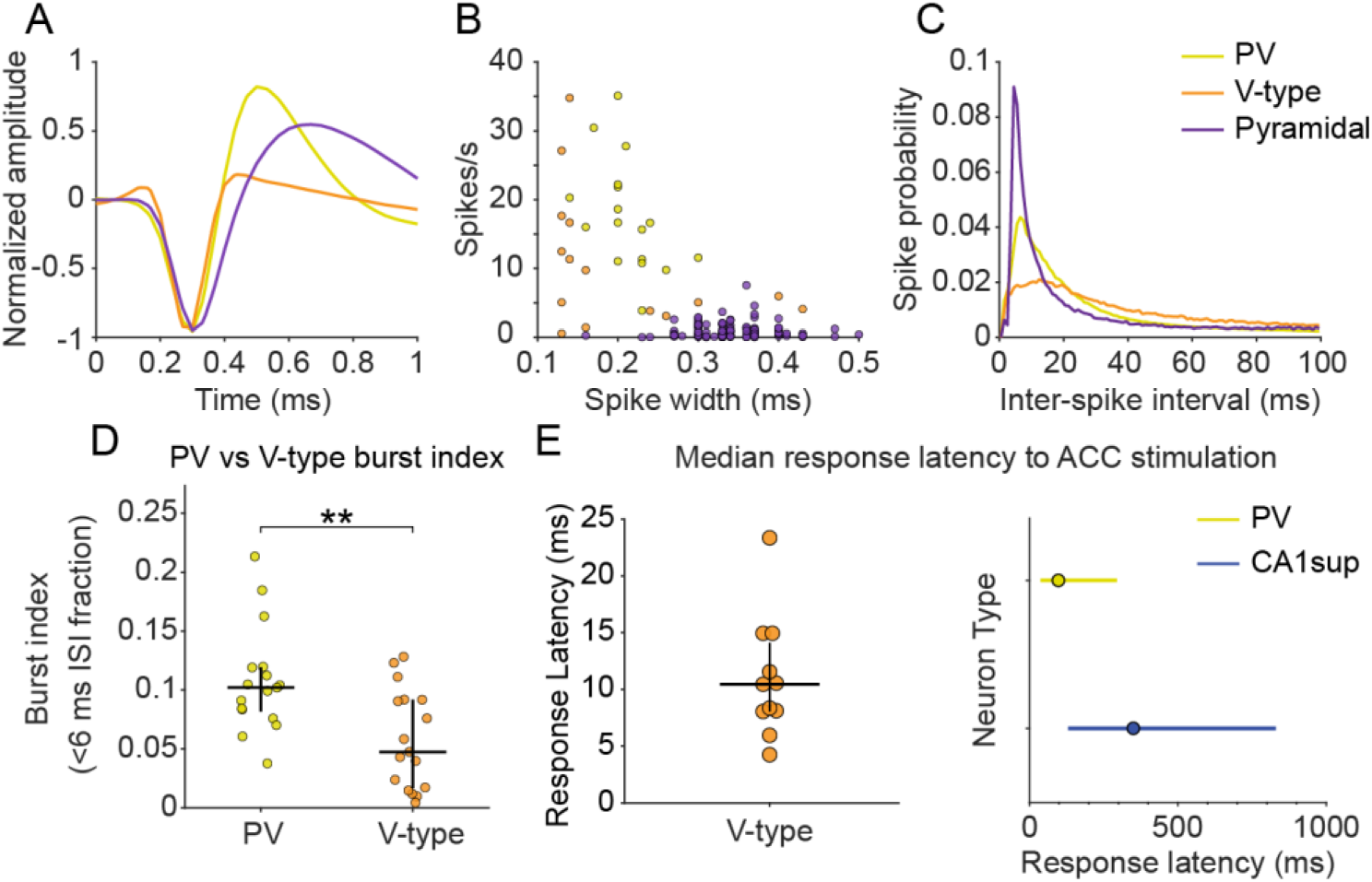
Putative Classifications of CA1 neurons. **A**, Normalized waveform averaged across all recorded neurons for each neuron subtype in optogenetic stimulation studies (Pyramidal N = 130, PVs N = 17, V-type N = 17). **B,** Putative classifications of neurons based on neuron firing rate and spike width. **C,** Inter-spike interval for each neuron subtype. **D**, PV and V-type interneuron burst-index comparison. A two-sided Mann-Whitney U test revealed that PV neurons (N = 17) showed a significantly higher burst index than V-type neurons (N = 17), p < .01. Points indicate individual neurons; horizontal black lines indicate group medians and vertical black lines indicate interquartile ranges. E, Median response latency of significantly responsive neurons following ACC stimulation. Left, V-type interneuron had a median latency 10.5 ms (N = 11, IQR = 8.1–13.3 ms). Points indicate individual neurons; horizontal black lines indicate group medians and vertical black lines indicate interquartile ranges. Right, PV interneurons had a median latency of 97.5ms (N = 17, IQR = 37.5–295 ms). CA1sup neurons had a median latency to respond of 370 ms (N = 54, IQR = 130–830 ms). Points indicate median and horizontal lines interquartile ranges (IQR).

**Figure 6—figure supplement 2.**
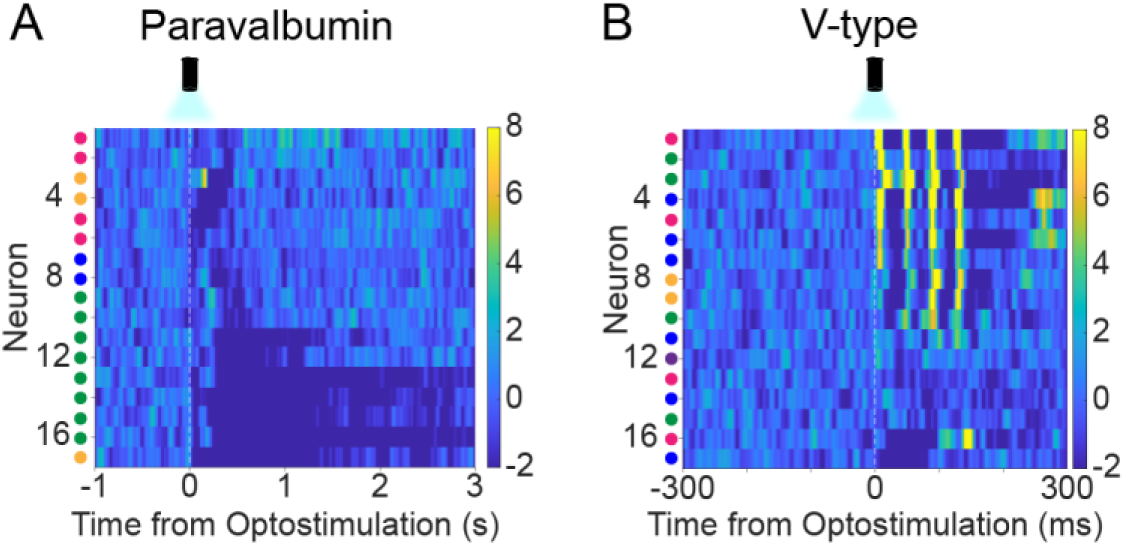
Per animal CA1 interneurons response to ACC stimulation, **A**, Heatmap of PV interneuron activity following stimulation of the ACC (bin, 5 ms; N = 17). Colored spheres to the left of each row indicate animal identity. Neurons from the same animal share the same color. **B,** Heatmap of V-type interneurons spiking activity following stimulation of the ACC (bin, 1 ms; N = 17).

